# Ongoing recombination in SARS-CoV-2 revealed through genealogical reconstruction

**DOI:** 10.1101/2021.01.21.427579

**Authors:** Anastasia Ignatieva, Jotun Hein, Paul A. Jenkins

**Affiliations:** Department of Statistics, University of Warwick, Coventry CV4 7AL, UK; Department of Statistics, University of Oxford, 24-29 St Giles’, Oxford OX1 3LB, UK; Department of Computer Science, University of Warwick, Coventry CV4 7AL, UK; The Alan Turing Institute, British Library, London NW1 2DB, UK

## Abstract

The evolutionary process of genetic recombination has the potential to rapidly change the properties of a viral pathogen, and its presence is a crucial factor to consider in the development of treatments and vaccines. It can also significantly affect the results of phylogenetic analyses and the inference of evolutionary rates. The detection of recombination from samples of sequencing data is a very challenging problem, and is further complicated for SARS-CoV-2 by its relatively slow accumulation of genetic diversity. The extent to which recombination is ongoing for SARS-CoV-2 is not yet resolved. To address this, we use a parsimony-based method to reconstruct possible genealogical histories for samples of SARS-CoV-2 sequences, which enables us to pinpoint specific recombination events that could have generated the data. We propose a statistical framework for disentangling the effects of recurrent mutation from recombination in the history of a sample, and hence provide a way of estimating the probability that ongoing recombination is present. We apply this to samples of sequencing data collected in England and South Africa, and find evidence of ongoing recombination.

## 1. Introduction

Ongoing mutation of the SARS-CoV-2 virus has received significant scientific and media attention since the start of the pandemic. The process of viral recombination has received far less coverage, but has the potential to have a drastic impact on the evolution of virulence, transmissibility, and evasion of host immunity (Simon-Loriere & Holmes, 2011). Recombination occurs when host cells are co-infected with different strains of the same virus, and during replication the genomes are reshuffled and combined before being packaged and released as new offspring virions, now potentially possessing very different pathogenic properties. This makes the presence of recombination a crucial factor to consider when developing vaccines and treatments. While the role of recombination between different coronaviruses in the emergence of SARS-CoV-2 has been widely studied, understanding its potential for ongoing recombination within human hosts has proved difficult.

The detection of ongoing recombination from a sample of genetic data is, in general, a very challenging problem. Only a fraction of recombination events significantly change the shape of a genealogy, and even then, mutations must occur on the correct branches of the genealogy in order to create patterns that are detectable in the data (Hein *et al*., 2004, Section 5.11). In evolutionary terms, a relatively short time period has passed since the start of the pandemic, so typical SARS-CoV-2 sequences differ only by a small number of mutations, meaning that recombination events are likely to be undetectable or leave only faint traces. Moreover, the effects of recombination can be indistinguishable from those of recurrent mutation (McVean *et al*., 2002), where mutations have occurred at the same site multiple times in the history of the sample. Coronaviruses are known to have relatively high recombination rates (Su *et al*., 2016), and cell culture studies indicate that this holds true for SARS-CoV-2 (Gribble *et al*., 2021). This suggests that ongoing intra-host recombination since the start of the pandemic should be commonplace, but detection efforts are thwarted by the slow accumulation of genetic diversity.

Early evidence of ongoing recombination in SARS-CoV-2 was presented by Yi (2020), who identified the presence of loops in reconstructed phylogenetic networks, which can arise as a consequence of recombination. A number of more recent reports have utilised methods based on classifying sequences into clades, and searching for those that appear to carry a mix of mutations characteristic to more than one clade. VanInsberghe et *al*. (2020) identified 1175 possible recombinants out of 537 000 analysed sequences; Varabyou et *al*. (2020) identified 225 possible recombinants out of 84 000; Jackson et *al*. (2021) have identified a small number of putative recombinants circulating in the UK. These methods are sensitive to the classification of sequences into clades, do not allow for the detection of intra-clade recombinants (thus underestimating the overall extent of recombination), and do not incorporate a framework for quantifying how likely it is that an observed pattern of incompatibilities has arisen through recombination rather than recurrent mutation. A number of studies have also failed to detect recombination signal, through the analysis of linkage disequilibrium and similar techniques (De Maio *et al*., 2020; van Dorp *et al*., 2020b; Nie *et al*., 2020; Tang *et al*., 2020; Wang *et al*., 2020; Richard *et al*., 2020). In general, a relatively small number of putative recombinant sequences have been identified to date, and there is a lack of compelling evidence for widespread recombination in SARS-CoV-2. Given the aforementioned causes for studies to be underpowered, the overall extent and importance of ongoing recombination for SARS-CoV-2 remains not fully resolved.

Phylogenetic analysis of SARS-CoV-2 data largely assumes the absence of recombination. Recombination can significantly influence the accuracy of phylogenetic inference (Posada & Crandall, 2002), distorting the branch lengths of inferred trees and making mutation rate heterogeneity appear stronger (Schierup & Hein, 2000). Moreover, when analysing data at the level of consensus sequences, the genealogy of a sample is related to the transmission network of the disease, with splits in the genealogy relating to the transmission of the virus between hosts. Models used for constructing genealogies and inferring evolutionary rates for this type of data cannot fully incorporate potentially important factors, such as geographical structure, patterns of social mixing, travel restrictions, and other non-pharmaceutical interventions, without making inference intractable. Relying on standard tree-based models can easily lead to biased estimates, with the extent of the error due to model misspecification being very difficult to quantify.

In this article, we use KwARG (Ignatieva *et al*., 2021), a parsimony-based method for reconstructing possible genealogical histories, to detect and examine crossover recombination events in samples of viral consensus sequences. This approach provides a concrete way of describing their genealogical relationships, sidestepping the challenges presented by discrepancies in clade assignment, enabling the detection of intra-clade recombination, avoiding the need to specify a particular model of evolution, and allowing for the explicit identification of possible recombination events in the history of a sample. Our method naturally handles both recombination and recurrent mutation, identifying a range of possible explicit genealogical histories for the dataset with varying proportions of both events types. Rather than using summary statistics calculated from the data, or focussing only on patterns of clade-defining SNPs, our method utilises all of the information contained in the patterns of incompatibilities observed in a sample, allowing for powerful detection and identification of possible recombinants. Moreover, we provide a nonparametric framework for evaluating the probability of a given number of recurrent mutations, thus quantifying how many recombinations are likely to have occurred in the history of a dataset. This allows for a more thorough and statistically principled assessment of the extent to which ongoing recombination is occurring.

We investigate the presence of ongoing recombination in SARS-CoV-2 using publicly available data from GISAID (Elbe & Buckland-Merrett, 2017), collected between November 2020 and February 2021. Using data from South Africa, we demonstrate that our method can detect recombination both when the sample contains sequences from multiple distinct lineages (‘inter-clade’), as well as all from the same lineage (‘intra-clade’). Further, we show that our method can accurately detect consensus sequences carrying patterns of mutations that are consistent with recombination, flagging these sequences for further investigation — and we demonstrate, using data from England, that it can identify both sequences arising as a result of sequencing errors due to sample contamination, aiding in identifying quality control issues, as well as sequences likely to be true recombinants. We validate our method using extensive simulation studies, and through application to MERS-CoV data, for which we find evidence of recombination, in agreement with previous studies.

## 2. Materials and Methods

### 2.1. SARS-CoV-2 data

Sequences were downloaded from GISAID, and aligned as described in SI Appendix, Section S1.1. Masking was applied to sites at the endpoint regions of the genomes, any multi-allelic sites, regions with many missing nucleotides in multiple sequences, and sites identified by De Maio *et al*. (2020) as being highly homoplasic or prone to sequencing errors. Strict quality criteria were applied, as detailed in SI Appendix, Section S1.2, to remove any sequences with a large number of ambiguous nucleotides, multiple non-ACTG characters, excessive gaps, and groups of clustered SNPs; additionally, sites identified by van Dorp *et al*. (2020a) as being prone to recurrent mutation were masked. These measures were aimed at reducing the possibility of including poor quality or contaminated sequences in the analysed samples, and also masking sites that are known to be highly homoplasic (either due to recurring sequencing errors, or due to the effects of selection).

Four samples were analysed: from South Africa, collected in (i) November 2020 (50 sequences, with 25 from lineage B.1.351, and 25 from other lineages) and (ii) February 2021 (38 sequences, all from lineage B.1.351), and from England, collected in (iii) November 2020 (80 sequences, with 40 sequences from lineage B.1.1.7 and 40 from other lineages within GISAID clade GR) and (iv) December 2020 – January 2021 (40 sequences within GISAID clade GR). Details of sample selection and processing are given in SI Appendix, Sections S1.3–S1.6.

### 2.2. Overview of methods

Our method consists of two main steps. Firstly, using KwARG, plausible genealogical histories are reconstructed for each sample, with varying proportions of posited recombination and recurrent mutations events. Then, using simulation, we approximate the distribution of the number of recurrent mutations that might be observed in a dataset of the same size as each sample. We use this to establish which of the identified genealogical histories is more plausible for the data at hand, and thus whether the presence of recombination events in the history of the given sample is likely.

This can be framed in the language of statistical hypothesis testing. The ‘null hypothesis’ is the absence of recombination. The test statistic *T* is the number of recurrent mutations in the history of the dataset; the null distribution of *T* is approximated through simulation. The observed value *T_obs_* is the minimal number of recurrent mutations required to explain the dataset in the absence of recombination, as estimated by KwARG. The ‘*p*-value’ is the probability of observing a number of recurrent mutations equal to or greater than *T_obs_*. Small *p*-values allow us to reject the null hypothesis, providing evidence that recombination has occurred. The reconstructed genealogies then allow for the detailed examination of possible recombination events in the history of the sampled sequences.

We emphasise that we make very conservative assumptions throughout, both in processing the data and in estimating the distribution of the number of recurrent mutations. Moreover, the number of recurrent mutations required to explain a given dataset computed by KwARG is (or is close to) a lower bound on the actual number of such events, and is likely to be an underestimate, making the reported *p*-values larger (more stringent).

### 2.3. Reconstruction of genealogies

The first step in our approach is to use a parsimony-based method to reconstruct possible genealogical histories for the given datasets.

#### 2.3.1. Incompatibilities in the data

Suppose that each site of the genome can mutate between exactly two possible states (thus excluding the possibility of triallelic sites, which we have masked from the data). Then the allele at each site can be denoted 0 or 1. If the commonly used *infinite sites* assumption is applied, at most one mutation can affect each site of the genome. The *four gamete* test (Hudson & Kaplan, 1985) can then detect the presence of recombination: if all four of the configurations 00, 01, 10, 11 are found in any two columns, then the data could not have been generated through replication and mutation alone, and at least one recombination event must have occurred between the two corresponding sites; the sites are then termed *incompatible*. If the infinite sites assumption is violated, the four gamete test no longer necessarily indicates the presence of recombination, as the incompatibilities could instead have been generated through recurrent mutation (McVean *et al*., 2002).

#### 2.3.2. Ancestral recombination graphs (ARGs)

All of the viral particles now in circulation had a common ancestor at the time of emergence of the virus, so sequences sampled at the present time can be united by a network of evolution going back to this shared ancestor through shared predecessors, termed the *ancestral recombination graph (ARG)* (Griffiths & Marjoram, 1997). As the sample consists of consensus sequences (at the level of one sequence per host), an edge of this network represents a viral lineage, possibly passing through multiple hosts before being sequenced at the present. An example of an ARG topology can be seen in Figure 4. Mutations are represented as points on the edges, labelled by the sites they affect. Considering the graph backwards in time (from the bottom up), the point at which two edges merge represents the time at which some sequences in the data coalesced, or have found a common ancestor. A point at which an edge splits into two corresponds to a recombination — the parts of the genome to the left and to the right of the breakpoint (whose site number is labelled inside the blue recombination node) are inherited from two different parent particles. The network thus fully encodes the evolutionary events in the history of a sample.

#### 2.3.3. Parsimonious reconstruction of histories

A sample of genetic sequences may have many possible histories, with many different corresponding ARGs. The *parsimony* approach to reconstructing ARGs given a sample of genetic data focuses on minimising the number of recombination and/or recurrent mutation events. This does not necessarily produce the most biologically plausible histories, but it does provide a lower bound on the number of events that must have occurred in the evolutionary pathway generating the sample. Thus, recombination can be detected in the history of a sample by considering whether the most plausible parsimonious solutions contain at least one recombination node.

Crucially, the parsimony approach does not require the assumption of a particular generative model for the data (such as the coalescent with exponential growth) beyond specifying the types of events that can occur. While this means that mutation and recombination *rates* cannot be inferred, it allows us to sidestep the need to specify a detailed model of population dynamics, which is particularly challenging for SARS-CoV-2 data. A parsimony-based approach is more appropriate when our focus is on interrogating the hypothesis that recombination is present at all. It also allows for the explicit reconstruction of possible events in the history of a sample, and thus allows us to identify recombinant sequences and uncover patterns consistent with the effects of sequencing errors.

#### 2.3.4. KwARG

KwARG (Ignatieva *et al*., 2021) is a program implementing a parsimony-based heuristic algorithm for reconstructing plausible ARGs for a given dataset. KwARG identifies ‘recombination only’ solutions (all incompatibilities are resolved through recombination events) and ‘recurrent mutation only’ solutions (all incompatibilities are resolved through additional mutation events), as well as interpolating between these two extremes and outputting solutions with a combination of both event types. KwARG allows for missing data and disregards insertions and deletions (we have deleted insertions from the alignment and treat deletions as missing data). KwARG seeks to minimise the number of posited recombination and recurrent mutation events in each solution, and the proportions of the two event types can be controlled by specifying ‘cost’ parameters. KwARG distinguishes between recurrent mutations that occur on the internal branches of the ARG from those can be placed on the terminal branches, which affect only one sequence in the input dataset, so can be examined separately for indications that they arose due to errors in the sequencing process.

KwARG was run on the data samples as detailed in SI Appendix, Section S3; an overview of the identified solutions is given in Tables 1a–1d.

**Table 1.** Summary of solutions identified by KwARG for each sample, and the probability of observing the corresponding number of recurrent mutations. First column: number of recombinations. Second column: number of recurrent mutations. Third column: probability of observing a number of recurrent mutations equal to that in the second column. Fourth column: corresponding *p*-values (probability of observing a number of recurrent mutations equal to or greater than that in the second column).

| (a) South Africa (Nov) |  |  |  | (b) South Africa (Feb) |  |  |  | (c) England (Jan) |  |  |  | (d) MERS-CoV |  |  |  |
| --- | --- | --- | --- | --- | --- | --- | --- | --- | --- | --- | --- | --- | --- | --- | --- |
| R | RM | $\mathbb{P}(RM)$ | $p$ | R | RM | $\mathbb{P}(RM)$ | $p$ | R | RM | $\mathbb{P}(RM)$ | $p$ | R | RM | $\mathbb{P}(RM)$ | $p$ |
| 10 | 0 | 0.28 | 1.00 | 7 | 0 | 0.52 | 1.00 | 10 | 0 | 0.11 | 1.00 | 9 | 0 | 0.42 | 1.00 |
| 8 | 1 | 0.35 | 0.72 | 5 | 1 | 0.34 | 0.48 | 8 | 1 | 0.24 | 0.89 | 7 | 1 | 0.36 | 0.58 |
| 6 | 2 | 0.23 | 0.37 | 3 | 2 | 0.11 | 0.14 | 6 | 2 | 0.27 | 0.65 | 6 | 2 | 0.16 | 0.22 |
| 4 | 3 | 0.10 | 0.14 | 2 | 3 | 0.03 | 0.03 | 4 | 3 | 0.20 | 0.38 | 5 | 3 | 0.05 | 0.06 |
| 3 | 4 | 0.03 | 0.04 | 1 | 4 | 0.00 | $5 \cdot 10^{-3}$ | 3 | 4 | 0.11 | 0.19 | 4 | 4 | 0.01 | 0.01 |
| 2 | 5 | 0.01 | 0.01 | 0 | 5 | 0.00 | $7 \cdot 10^{-4}$ | 2 | 5 | 0.05 | 0.08 | 3 | 5 | 0.00 | $2 \cdot 10^{-3}$ |
| 1 | 7 | 0.00 | $4 \cdot 10^{-4}$ | | | | | 1 | 6 | 0.02 | 0.03 | 2 | 10 | 0.00 | $< 1 \cdot 10^{-6}$ |
| 0 | 9 | 0.00 | $7 \cdot 10^{-6}$ | | | | | 0 | 14 | 0.00 | $1 \cdot 10^{-6}$ | 1 | 12 | 0.00 | $< 1 \cdot 10^{-6}$ |
| | | | | | | | | | | | | 0 | 16 | 0.00 | $< 1 \cdot 10^{-6}$ |

### 2.4. Evaluation of solutions

The next step in our approach is to determine which of the solutions identified by KwARG is more likely, by calculating the probability of observing the given number of recurrent mutations. To avoid making model-based assumptions on the genealogy of the sample, we use a nonparametric method inspired by the *homoplasy test* of Maynard Smith & Smith (1998).

The homoplasy test estimates the probability of observing the minimal number of recurrent mutations required to generate the sample in the absence of recombination, i.e. if the shape of the genealogy is constrained to be a tree. If this probability is very small then it provides evidence for the presence of recombination. The method is particularly powerful when the level of divergence between sequences is very low, as is the case with SARS-CoV-2 data, although it appears prone to false positives in the presence of severe mutation rate heterogeneity along the genome (Posada & Crandall, 2001). We calculate an empirical estimate 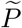 of mutation density along the genome from SARS-CoV-2 data, which does not suggest the presence of extreme heterogeneity, and then use this estimate to simulate the distribution of the number of recurrent mutations that are observed in a sample.

The *i*-th entry of the vector 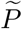, for *i* ∈ {1,…, 29 903}, gives an estimated probability that when a mutation occurs, it affects the *i*-th site of the genome. Details of our method for estimating 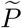 are presented in SI Appendix, Section S4. Briefly, this estimate is calculated by examining the locations of sites that have undergone at least one mutation (segregating sites) using GISAID data collected in February 2021. If the mutation rate were constant along the genome, we would expect segregating sites to be spread uniformly throughout the genome; uneven clustering of the mutations gives an indication of mutation rate heterogeneity. We use a nonparametric method (wavelet decomposition) to estimate 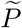 from the observed positions of segregating sites, taking into account the dependence of the mutation rate on the base type of the nucleotide undergoing mutation, which is significant for SARS-CoV-2 (Simmonds, 2020; Koyama *et al*., 2020). The resulting estimate is shown in Figure 1.

**Figure 1.**
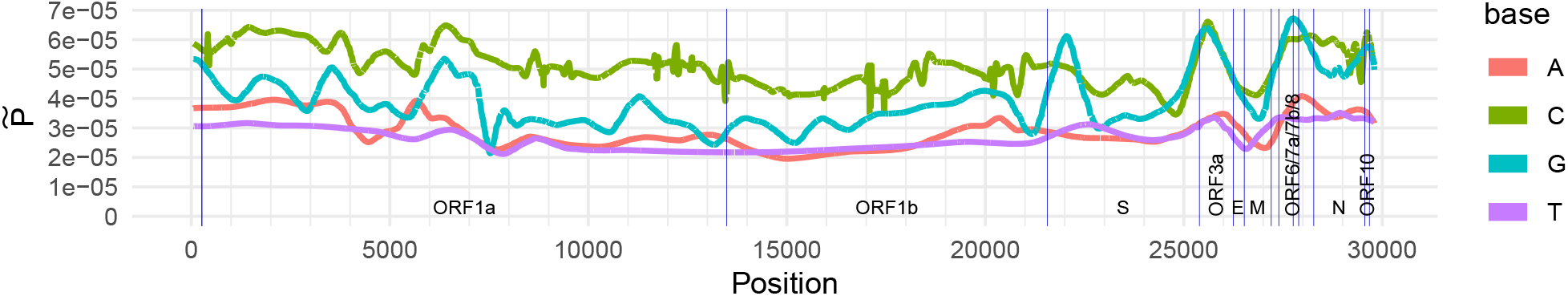
Estimate 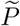 of the probability of a mutation falling on each site of the SARS-CoV-2 genome. Blue vertical lines mark endpoints of the labelled ORFs and genes as per Wu *et al*. (2020).

The estimate of 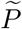 is then used to approximate the distribution of the number of recurrent mutations observed in a sample, using a simulation approach. We simulate the process of mutations falling along the genome until the simulated number of segregating sites matches that observed in the sample; the vector 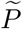 controls where on the genome each mutation falls. The number of recurrent mutations (instances where mutations fall on the same site multiple times) is recorded. This procedure is repeated for 1 000 000 iterations and a histogram of the results is constructed. The resulting probabilities and corresponding *p*-values are shown in the third and fourth columns of Tables 1a-1d.

## 3. Results

### 3.1. Validation on simulated data

#### 3.1.1. False positives

The accuracy of the presented method depends on an assumption that there are no highly homoplasic sites (arising either due to selection or repeated sequencing errors) that have not been masked. If this assumption were violated, the estimate 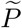 would be missing ‘spikes’ of high probability at the corresponding positions, biasing the simulated null distribution to underestimate the number of recurrent mutations, and potentially leading to false positive results.

We investigated the validity of this assumption through simulation as described in SI Appendix, Section S4.3.2, by inflating the mutation probability of a subset of 0 to 200 sites in the vector 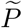 by a factor *H*, simulating data with the resulting mutation rate map in the absence of recombination with parameters that appear reasonable for SARS-CoV-2, and checking whether our method would (incorrectly) reject the null hypothesis. The results are presented in the left panel of Figure 2.

**Figure 2.**
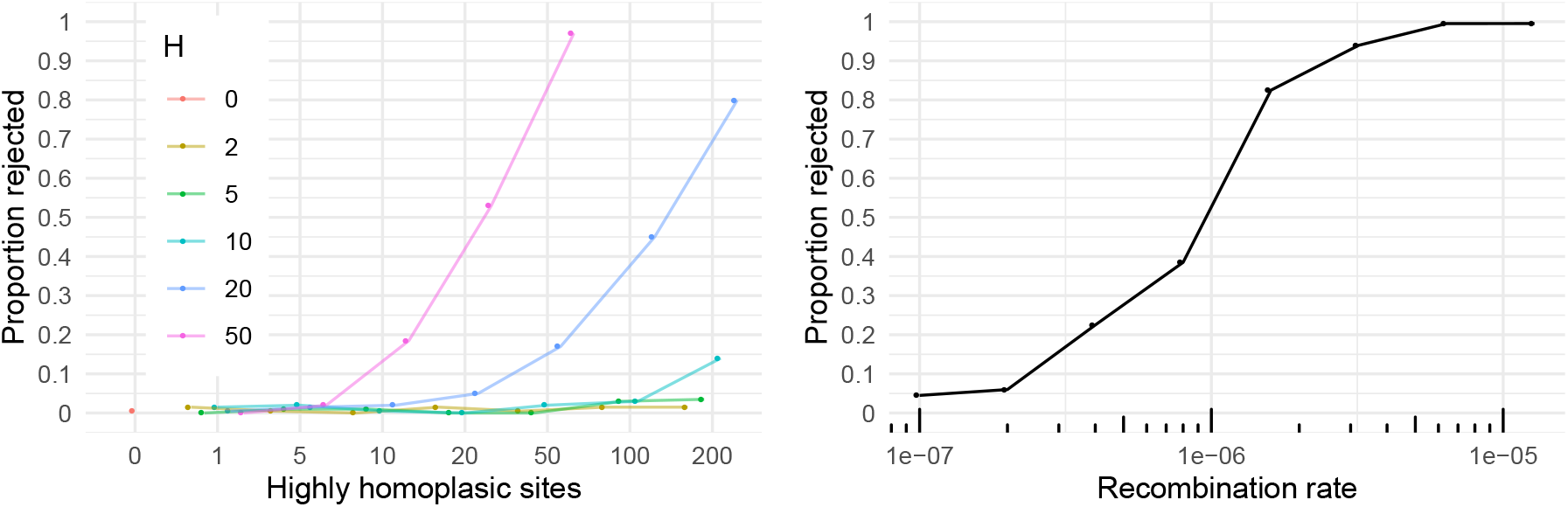
Left panel: *x*-axis shows number of added highly homoplasic sites, with the corresponding entries of 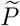 multiplied by the factor *H* (colours); *y*-axis shows the proportion of simulated datasets (out of 200 for each combination of parameters) for which the null hypothesis was (incorrectly) rejected with *p* < 0.05. Right panel: *x*-axis shows recombination rate (per site per generation) used to simulate 200 datasets, *y*-axis shows proportion of datasets for which the null hypothesis was rejected with *p* < 0.05.

False positives were seen in only 0.5% of cases when there are no highly homoplasic sites, demonstrating that our method conservatively overestimates the computed *p*-values. The proportion of false positives only increases significantly when a large number of extremely homoplasic sites is present, showing that our method is reasonably robust to violations of this assumption. Particularly as we apply a conservative strategy in masking sites known to be homoplasic, this shows that our method is unlikely to falsely indicate the presence of recombination.

#### 3.1.2. Detectable recombination rate

The power of our test in detecting the presence of recombination was investigated for a range of recombination rates *ρ*, by simulating datasets as described in SI Appendix, Section S4.3.3, and recording how often the null hypothesis of no recombination could be rejected (with *p* < 0.05). The results are shown in the right panel of Figure 2, demonstrating that this occurred in 4.5% of cases for *ρ* = 1 · 10^-7^ per site per generation, rising to 99.5% of cases for *ρ* = 1 · 10^-5^. The simulations were performed using parameters that appear reasonable for SARS-CoV-2; the results suggest that our method is sufficiently powerful for detecting recombination if the recombination rate is higher than around *ρ* = 1 · 10^-6^ ≈ 4 · 10^-5^ per site per year.

### 3.2. Identification of recombinant sequences

All sequences collected in England in December 2020 – January 2021, labelled as belonging to clade GR in GISAID, were downloaded and processed as described in SI Appendix, Section S1.6. The resulting sample comprises 40 sequences with 276 variable sites.

An illustration of the sample is provided in SI Appendix, Figure S4. Choosing a solution with no recombinations, the sites of fourteen recurrent mutations identified by KwARG are highlighted with red (yellow) crosses, where the recurrent mutations fall on the terminal (internal) branches of the ARG. The sequencing protocol used by the COVID-19 Genomics UK Consortium, the submitters of the data, generates short amplicons of under 400 bp in length, and none of the identified sites of recurrent mutations fall into the same amplicon region, making it less likely that the results are due to sample contamination or other sequencing artifacts. The probability of observing the required *T_obs_* = 14 or more recurrent mutations is *p* = 1 · 10^-6^, which strongly indicates the presence of recombination.

Considering the results in Table 1c, three recurrent mutations can have the same effect as six of the identified recombination events (compare row (*R,RM*) = (10,0) with (*R,RM*) = (4,3)), suggesting that recurrent mutation offers a more parsimonious explanation for at least part of the patterns seen in the data. One of these recurrent mutations consistently occurs at site 22 227; the other two can be placed either at the same site 9 693, or at sites 9 693 and 12 067. The probability of observing five or fewer recurrent mutations is 0.97, which suggests that, with high probability, at least two recombination have occurred in the history of the sample. An example of an ARG with two recombination events is shown in Figure 3.

**Figure 3.**
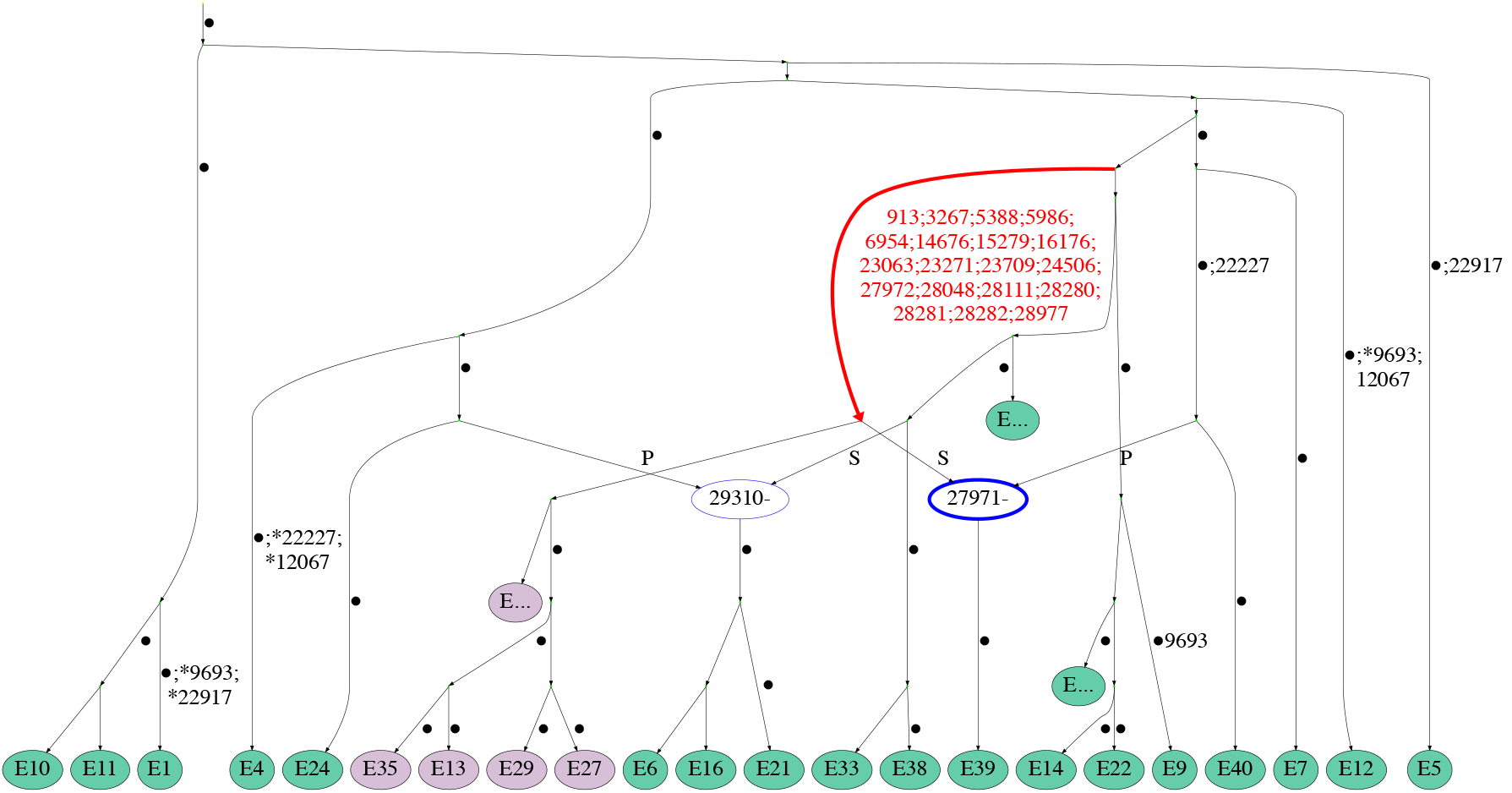
Example of an ARG for the England (January) dataset. Recombination nodes are shown in blue, labelled with the recombination breakpoint, with the offspring sequence inheriting part of the genome to the left (right) of the breakpoint from the parent labelled “P” (“S”). Recurrent mutations are prefixed with an asterisk. Edge carrying the characteristic mutations of lineage B.1.1.7 is highlighted in red; nodes corresponding to sequences from lineage B.1.1.7 are coloured purple. For ease of viewing, some parts of the ARG have been collapsed into nodes labelled “E…”. Edges are labelled by positions of mutations (some mutated sites are not explicitly labelled and are denoted by a dot instead).

It is striking that eight of the recurrent mutations seen in Figure S4 can be placed in the same sequence E39. Indeed, Figure 3 shows that the corresponding incompatibilities in the data can be resolved by just one recombination event between sequence E40 and a sequence from lineage B.1.1.7; the corresponding recombination node is shown in bold. The sequence E39 has previously been identified as a possible recombinant by Jackson *et al*. (2021), demonstrating that our method can clearly highlight mosaic sequences in addition to quantifying the probability that recombination has occurred in the history of the dataset.

### 3.3. Detection of intra-clade recombination

All sequences collected in South Africa in February 2021 were downloaded and processed as described in SI Appendix, Section S1.4. The resulting sample comprises 38 sequences with 151 variable sites, all from the same lineage B.1.351.

Initial examination of the solutions identified by KwARG show that at least 8 recurrent mutations are required to construct a valid ARG for this sample in the absence of recombination. However, it was noted that three of these recurrent mutations occur at the same site 28 254. This may imply that the site is highly mutable, which could be due to repeated sequencing errors, or as a consequence of selection. We note that this demonstrates the usefulness of our approach in identifying potentially highly homoplasic sites.

This position was masked from the sample before re-running the analysis. The probability of observing the re-calculated value of *T_obs_* = 5 or more recurrent mutations is *p* = 7 · 10^-4^, strongly suggesting the presence of recombination. The probability of observing two or fewer recurrent mutations is 0.97, which indicates that with high probability, at least three recombination events have occurred in the history of the dataset.

### 3.4. Detection of inter-clade recombination

All sequences collected in South Africa in November 2020 were downloaded and processed as described in SI Appendix, Section S1.3, to create a sample of 50 sequences with 207 variable sites, with 25 belonging to lineage B.1.351 (labelled SAN1-SAN25), and 25 to other lineages (labelled SAO1-SAO25).

An initial run of KwARG demonstrated that, notably, one recurrent mutation occurs at site 28 254, further suggesting that this site is excessively prone to recurrent mutation. This site was therefore masked before re-running the analysis. An illustration of the sample is provided in SI Appendix, Figure S5. The sites of nine recurrent mutations identified by KwARG are highlighted with red crosses (choosing a solution with no recombinations, and where the recurrent mutations fall on the terminal branches of the ARG). The probability of observing the required *T_obs_* = 9 or more recurrent mutations is *p* = 7 · 10^-6^, strongly suggesting the presence of recombination.

The probability of observing three or fewer recurrent mutations is 0.96, which indicates that, with high probability, at least four recombination events have occurred in the history of the dataset. Indeed, Table 1 shows that three recurrent mutations can remove the necessity of six recombination events, suggesting that recurrent mutation offers a more parsimonious explanation than recombination for the remaining incompatibilities in the data. Examination of the KwARG solutions shows that these recurrent mutations consistently occur at sites 4 093, 11 230, and 25 273. An ARG with recurrent mutations at these three sites is shown in Figure 4; edges carrying the characteristic mutations of lineage B.1.351 are highlighted in red.

**Figure 4.**
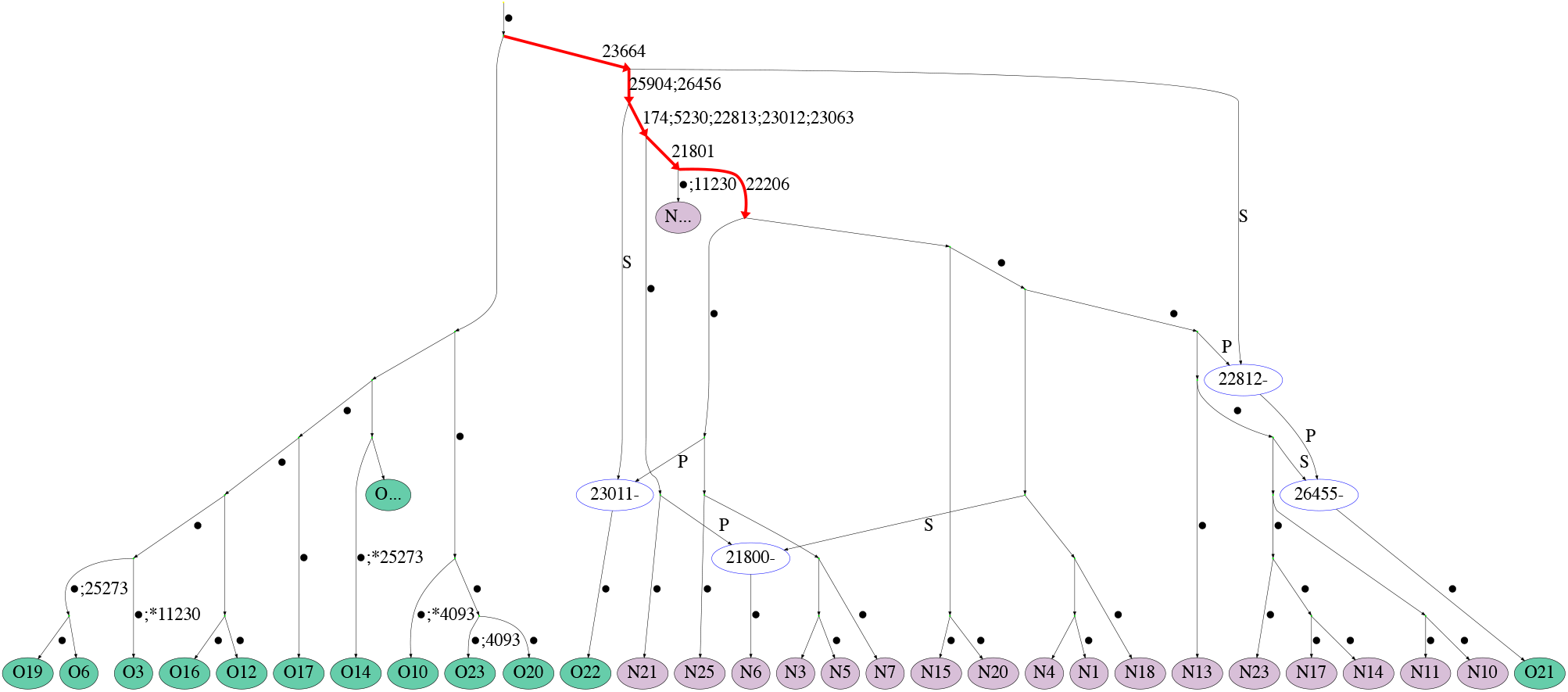
Example of an ARG for the South Africa (November) dataset (the “SA” prefix of each sequence reference number is dropped for ease of viewing). Recombination nodes are shown in blue, labelled with the recombination breakpoint, with the offspring sequence inheriting part of the genome to the left (right) of the breakpoint from the parent labelled “P” (“S”). Recurrent mutations are prefixed with an asterisk. For ease of viewing, some parts of the ARG have been collapsed into nodes labelled “O…” and “N…” (containing sequences labelled SAO and SAN, respectively). Edges are labelled by positions of mutations (some mutated sites are not explicitly labelled and are denoted by a dot instead).

The sequences SAO21 and SAO22 carry three and two of the identified nine recurrent mutations, respectively, when recombination is prohibited in reconstructing the genealogy. Both of these sequences carry some of the mutations characteristic of lineage B.1.351; this is demonstrated in Figure 5, where the two sequences are compared to two other typical sequences from lineage B.1.351. Examination of the KwARG solutions shows that a recombination in Sequence SAO21 just after site 22 812 has the same effect as the recurrent mutations at sites 22 813 and 23 012, and a recombination in Sequence SAO22 just after site 23 011 has the same effect as the recurrent mutations at sites 23 012 and 23 063. This suggests that the patterns of incompatibilities observed in these two sequences are consistent with recombination; a possible sequence of recombination events generating these sequences can be seen in the ARG in Figure 4.

**Figure 5.**
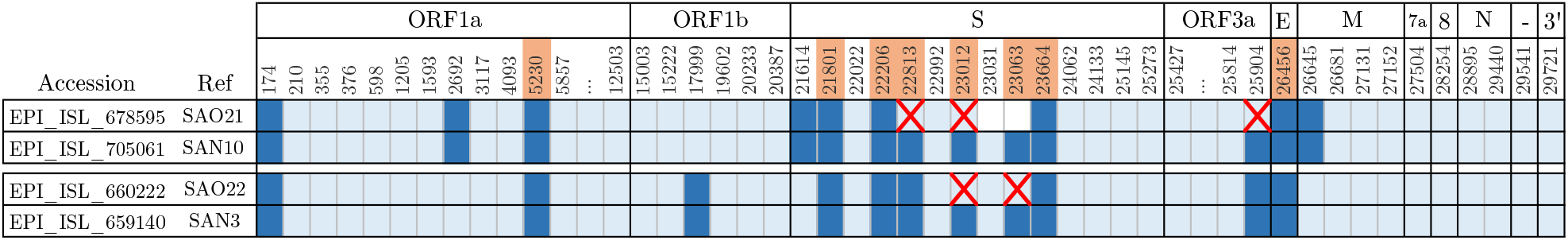
Comparison of sequences SAO21, SAO22 and the characteristic mutations for lineage B.1.351. Columns correspond to positions along the genome; uninformative sites (with all 0’s or 1’s) and those with singleton mutations (with exactly one 1) are not shown. Light blue: ancestral state, dark blue: mutated state, white: missing data. Red crosses highlight sites of recurrent mutations identified by KwARG. Sites bearing the characteristic (non-synonymous) mutations of lineage B.1.351 (Tegally *et al*., 2020) are highlighted in orange.

### 3.5. Identification of sequencing errors due to cross-contamination

All sequences labelled as GISAID clade GR, collected in England in November 2020, were aligned, masked, and processed as detailed in SI Appendix, Section S1.5. The quality criteria detailed in SI Appendix, Section S1.2 were *not* applied in this case. The resulting sample comprises 80 sequences with 363 variable sites, 40 of which belong to lineage B.1.1.7 (labelled EN1-EN40) and 40 to other lineages (labelled EO1-EO40).

The results showed that in the absence of recombination, at least 15 recurrent mutations were required to explain the incompatibilities observed in this sample. However, it was identified that six of these recurrent mutations could be placed in the same sequence EO40, as illustrated in Figure 6. The sequence EO40 appeared to carry some of the mutations carried by sequence EO32, and some of the mutations characteristic of lineage B.1.1.7, strongly suggesting that this sequence was a recombinant.

**Figure 6.**
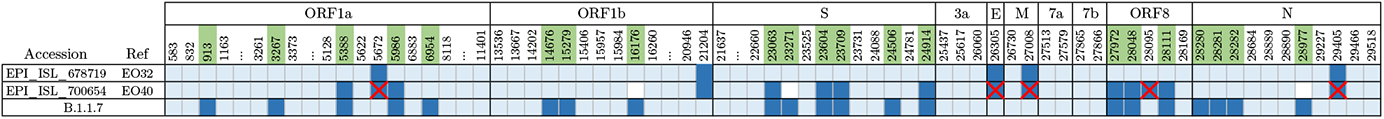
Comparison of sequences EO32, EO40 and the characteristic mutations of lineage B.1.1.7. Columns correspond to positions along the genome; uninformative sites (with all 0’s or 1’s) and those with singleton mutations (with exactly one 1) are not shown. Light blue: ancestral state, dark blue: mutated state, white: missing data. Red crosses highlight locations of the recurrent mutations identified by KwARG. Sites bearing the characteristic mutations of lineage B.1.1.7 (Rambaut *et al*., 2020) are highlighted in green.

Our findings prompted further investigation by the submitters of this sequence, which revealed the signal to be the result of significant contamination of the genetic sample causing multiple errors in the consensus sequence, rather than a result of intra-host recombination. The sequence has subsequently been removed from GISAID.

### 3.6. Recombination detection for MERS-CoV data

MERS-CoV sequences collected in Saudi Arabia in January-March 2015 were downloaded from the NCBI virus database (Hatcher *et al*., 2017), and aligned, masked, and processed as described in SI Appendix, Section S2. The resulting sample consists of 19 sequences with 197 variable sites.

The dataset is illustrated in SI Appendix, Figure S6. The locations of recurrent mutations identified by KwARG are shows as red and yellow crosses, corresponding to recurrent mutations occurring on the terminal and internal branches of the ARG, respectively. In the absence of recombination, at least *T_obs_* = 16 recurrent mutations are required, which has probability *p* < 1 · 10^−6^, strongly suggesting the presence of recombination. The probability of observing three or fewer recurrent mutations is 0.99, suggesting that at least five recombinations have occurred in the history of the sample. An ARG with five recombination nodes, showing a possible history of the dataset, is shown in SI Appendix, Figure S7.

A group of four identical sequences (M16-M19, shown in purple in Figure S7) appear to carry a characteristic set of shared mutations that strongly differentiates them from the other sequences in the sample. Five of the identified recurrent mutations affect this group, occurring in a relatively short stretch of the genome, suggesting that these patterns are indicative of recombination with other sequences in the sample carrying these mutations.

Five of the other identified recurrent mutations can be placed in one sequence (M11), which appears to carry a mixture of mutations from the group identified above and other sequences in the sample, which is consistent with recombination. This sequence does not match any others in the dataset, so it is possible that this is the result of sequencing errors or sample contamination. If this sequence is removed from the sample, at least *T_obs_* = 9 recurrent mutations are still required to explain the observed incompatibilities, which has probability *p* < 1 · 10^-6^, still strongly suggesting that recombination is present. This agrees with previous reports of within-host recombination for MERS-CoV (Zhang *et al*., 2016; Dudas & Rambaut, 2016; Sabir *et al*., 2016).

## 4. Discussion

The method presented in this article offers a clear and principled framework for recombination detection, which can be interpreted as a hypothesis testing approach. We make very conservative assumptions throughout, demonstrating on both real and simulated data that the method achieves a very low rate of false positive results, while offering powerful detection of recombination at even relatively low values of recombination rate. We use nonparametric techniques at each stage, to avoid making assumptions on the process generating the data, and thus circumvent issues with model misspecification. Our method allows us to gain clear insights into the evolutionary events that may have generated the given sequences, offering easily interpretable results. The method detects sequences carrying patterns consistent with recombination, demonstrating its effectiveness as a tool for flagging sequences with distinctive patterns of incompatibilities for further detailed investigation.

Our results clearly indicate the presence of recombination in the history of the analysed SARS-CoV-2 sequencing data, suggesting a recombination rate greater than around 4 · 10^-4^ per site per year. One of the main limitations of our method is that KwARG does not scale well to large datasets. However, while studies relying on clade assignment and statistics such as linkage disequilibrium have identified that recombination occurs at very low levels (VanInsberghe *et al*., 2020; Varabyou *et al*., 2020) or is unlikely to be occurring at a detectable level (De Maio *et al*., 2020; van Dorp *et al*., 2020b; Nie *et al*., 2020; Tang *et al*., 2020; Wang *et al*., 2020; Richard *et al*., 2020) even when analysing vast quantities of sequencing data, our method is powerful enough to detect the presence of recombination using even relatively small samples. Moreover, the testing framework could potentially be used in combination with other methods for reconstructing ARGs, including ones not relying on the parsimony assumption, with appropriate modifications to control the false positive rate and ensure validity of the results.

Recombination can occur when the same host is co-infected by two different strains, which has been noted to occur in COVID-19 patients (Samoilov *et al*., 2020), and could become more likely with the emergence of more transmissible variants. We note that the potential mosaic sequences we identified in the South Africa sample from November are represented only once in the data. This could be due to a lack of onward transmission, as recombinants are likely to reach a detectable level at a relatively late stage in the infection cycle. It could also indicate that the sequences arose due to either contamination of the sample during processing, or the misassembly of two distinct (non-recombinant) strains present in the same sample, as was identified to be the case for one sequence in the England sample from November.

We also note that while we sought to mask any sites known to be highly homoplasic, we cannot rule out that some of the identified recurrent mutations did arise multiple times as a consequence of selection or as a result of repeated sequencing errors. However, we have demonstrated that the solutions presented by KwARG can be examined for the presence of highly mutable sites, and have identified using both samples from South Africa that this appears to be the case for site 28 254 (located proximal to the stop codon of ORF8).

Our findings suggest that care should be taken when performing and interpreting the results of analysis based on the construction of phylogenetic trees for SARS-CoV-2 data. The presence of recombination, as well as other factors complicating the structure of the transmission network of the virus, strongly suggests that tree-based models are not appropriate for modelling SARS-CoV-2 genealogies, and inference of evolutionary rates based on such methods may suffer from errors due to model misspecification that are difficult to quantify.

Due to the high level of homogeneity between sequences, the effects of recombination will be either undetectable or indistinguishable from recurrent mutation in the majority of cases. However, as genetic diversity builds up over longer timescales, the effects of recombination may become more pronounced. Particularly in light of the recent emergence of new variants, the rapid evolution of the virus through recombination between strains with different pathogenic properties is a crucial risk factor to consider. This highlights the need for continuous monitoring of the sequenced genomes for new variants, to enable the early detection of novel recombinant genotypes, and for further work on the quantification of recombination rates and identification of recombination hotspots along the genome.

## 5. Data and code availability

SARS-CoV-2 sequencing data is publicly available from GISAID at gisaid.org upon free registration. MERS-CoV data is publicly available from the NCBI Virus database at ncbi.nlm.nih.gov/labs/virus. Code used in processing the data and carrying out the analysis is available at github.com/a-ignatieva/sars-cov-2-recombination.

## 6. Acknowledgements

We thank the maintainers and contributors of the GISAID database; a full table of acknowledgements for the data used is provided at github.com/a-ignatieva/sars-cov-2-recombination/GISAID_acknowledgements. We thank Oliver Pybus for help with identifying quality issues affecting sequence EO40. This work was supported by the EPSRC and MRC OxWaSP Centre for Doctoral Training (EPSRC grant EP/L016710/1), and by the Alan Turing Institute (EPSRC grant EP/N510129/1).

## Supporting Information

### S1. Data: SARS-CoV-2

#### S1.1. Alignment and masking

SARS-CoV-2 sequences were downloaded from GISAID (Elbe & Buckland-Merrett, 2017), filtering for those labelled as complete (>29 000bp), collected from human hosts, and excluding any with more than 5% ambiguous nucleotides and incomplete collection dates. Although SARS-CoV-2 is an RNA virus, we refer to nucleotides by their DNA type for consistency with the sequencing data (i.e. the base type T corresponds to U on the actual SARS-CoV-2 genome).

Alignment to the reference sequence collected in Wuhan in December 2019 (Wu *et al*., 2020) (GISAID accession: EPI_ISL_402125, GenBank: MN908947.3) was performed using MAFFT v7.475 (Katoh & Standley, 2013), with the options: `auto, keeplength, preservecase, addfragments`.

The following sites were masked from the data:

- the endpoint regions with a large number of missing nucleotides (1-55bp and 29 804-29 903bp);
- 322 further sites identified as problematic by De Maio *et al*. (2020) (prone to sequencing errors, known to be excessively homoplasic, or otherwise of questionable quality);
- any multi-allelic sites.

#### S1.2. Quality criteria

Any sequences failing the following quality criteria were removed:

- at most 500 missing nucleotides (excluding start and end of alignment);
- at most 1 non-ACTG character;
- at most 25 gaps;
- no SNP clusters (more than 6 SNPs in a window of 100 nucleotides, excluding known clusters of mutations).

Nextclade (Hadfield *et al*., 2018, tool available at clades.nextstrain.org) was used to check sampled sequences against these criteria (and it was ensured that any sequences assigned a score of “bad” by the tool were removed). In addition, the 198 sites identified by van Dorp *et al*. (2020, Supplementary Table S5) as potentially highly homoplasic were masked.

#### S1.3. South Africa (November)

All sequences collected in South Africa in November 2020 were downloaded and aligned as described in Section S1.1. Removing 48 sequences flagged by the submitter as containing long stretches of ambiguous nucleotides, and applying the quality criteria in Section S1.2, left a total of 278 sequences.

The aligned sequences were split into the datasets *SA_N_* (the 177 sequences labelled as belonging to variant 501Y.V2 in GISAID) and *SA_O_* (the other 101 sequences). A sample of 25 sequences from each of *SA_O_* and *SA_N_* was selected at random using SeqKit (Shen *et al*., 2016).

Masking was carried out as described in Section S1.1; in addition, sites 22 266–22 745 were masked, as many of the sequences contained a large number of ambiguous nucleotides at these positions. No further multi-allelic sites were identified. Of the total 1125 masked positions, 28 corresponded to segregating sites in the dataset.

The resulting sample comprises 50 sequences with 207 variable sites. The corresponding GISAID accession numbers and collection dates are given in Table S1.

#### S1.4. South Africa (February)

All sequences collected in South Africa in February 2021 were downloaded, aligned, and masked as described in Section S1.1, also masking sites 22 266-22 745; no additional multi-allelic sites were identified. The quality filters detailed in Section S1.2 were applied. One sequence in the resulting sample was not from lineage B.1.351 and was removed. Of the total 1 125 masked positions, 17 corresponded to segregating sites.

The resulting sample consists of 38 sequences, all from lineage B.1.351, with 151 variable sites. The corresponding GISAID accession numbers and collection dates are given in Table S2.

#### S1.5. England (November)

All sequences labelled as clade GR, collected in England in November 2020, were downloaded and aligned as per Section S1.1. Exact duplicates of sequences in the dataset were removed, to avoid including identical sequences in the sample. The sequences were then split into datasets *E_N_* (934 sequences labelled as belonging to lineage B.1.1.7) and *E_O_* (the other 2 650 sequences).

A sample of 40 sequences from each of *E_O_* and *E_N_* was then selected at random using SeqKit. Sites were masked as detailed in Section S1.1. Three multi-allelic sites were identified and masked, at positions 12 067, 21 724, and 22 992. Of the total 477 masked positions, 10 corresponded to segregating sites in the dataset. The quality control criteria in Section S1.2 were *not* applied to this sample.

The resulting sample comprises 80 sequences with 363 variable sites. The corresponding GISAID accession numbers and collection dates are given in Table S3.

#### S1.6. England (January)

All sequences labelled as clade GR, collected in England in December 2020 – January 2021, were downloaded and aligned as per Section S1.1. Sites were masked as detailed in Section S1.1, and the quality filters detailed in Section S1.2 were applied. A sample of 38 sequences was selected at random using SeqKit, from among sequences uploaded by the COG UK Consortium; additionally, the sequence EPI_ISL_994038 (E39) identified as a potential recombinant by Jackson *et al*. (2021), and its potential parent sequence EPI_ISL_820233 (E40), were included. Five multi-allelic sites were identified and masked, at positions 21255, 23 604, 24 914, 28 310, and 29 227. Of the total 660 masked positions, 35 corresponded to segregating sites in the dataset.

The resulting sample comprises 40 sequences with 276 variable sites. The corresponding GISAID accession numbers and collection dates are given in Table S4.

### S2. Data: MERS-CoV

MERS-CoV sequences were downloaded from the NCBI Virus database (Hatcher *et al*., 2017), filtering for those labelled as complete, human host, collected in Saudi Arabia in January-March 2015. Alignment to the reference sequence (HCoV-EMC/2012, accession number NC_019843.3) was performed using MAFFT, with the same options as in Section S1.1. Masking of the first and last 150 sites of the alignment was performed. Of the 300 masked sites, two were segregating in the dataset; no multi-allelic sites were identified. The resulting sample consists of 19 sequences with 197 variable sites. The corresponding accession numbers are given in Table S5.

### S3. KwARG

KwARG seeks to minimise the number of posited recombination and recurrent mutation events in each solution, and the proportions of the two event types can be controlled by specifying input ‘cost’ parameters *C_SE_, C_RM_, C_R_*, and *C_RR_*, corresponding to penalties assigned to recurrent mutations on the terminal branches of the ARG, those on internal branches, recombination events, and two consecutive recombination events (which can mimic the effects of gene conversion), respectively. For instance, setting (*C_SE_, C_RM_, C_R_, C_RR_*) = (0.5, 0.51, 1.0, 2.0) is likely to produce solutions with more recurrent mutations than recombinations, as the cost of recurrent mutations is lower, favouring placing recurrent mutations on the terminal branches of the ARG where possible. Recurrent mutations on the terminal branches of the ARG affect only one sequence in the input dataset, so can be examined separately for indications that they arose due to errors in the sequencing process.

KwARG implements a method of randomly exploring the space of possible ARGs, so it should be run multiple times for each configuration of input parameters, and the best identified solutions (with the minimal number of posited recombinations and/or recurrent mutations) then selected for analysis. An input parameter T (the ‘annealing temperature’) controls the extent of this random exploration.

#### S3.1. Parameter settings

For each dataset, KwARG was run *Q* = 500 times for each combination of the following values of the annealing parameter *T* and event costs (*C_SE_, C_RM_, C_R_, C_RR_*):

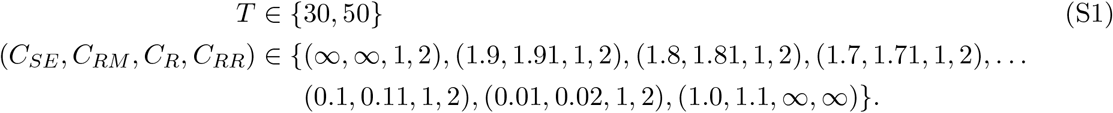

For MERS-CoV, the root was left unspecified. For SARS-CoV-2, the reference sequence used for alignment was set as the root. This reference sequence is a genome collected in Wuhan in December 2019 (Wu *et al*., 2020), giving the most likely rooting based on the available epidemiological evidence; our results do not change significantly if the root is left unspecified.

### S4. Null distribution for SARS-CoV-2

#### S4.1. Distribution of the number of recurrent mutations

Let *M* be the length of the genome, and let *m* be the number of observed variable sites in the sample. We are interested in estimating the distribution of the number of recurrent mutations that have occurred; that is, the excess number of mutation events beyond the minimum *m* needed to explain the variability in the sample.

Regardless of any modelling assumptions on the evolution of a given sample or the genealogical relationships between the sequences, it is clear that at least *m* mutation or sequencing error events must have occurred in the history of the sample (here, a ‘sequencing error’ refers to the variant at a site being incorrectly called during the sequencing process). Suppose that each time such an event occurs (disregarding which particular sequence is affected), a position on the genome is selected at random with replacement, according to a probability vector *P* of length M. This corresponds to assuming that (i) such events occur independently from each other, (ii) all sequences have the same probabilities *P* of a mutation or sequencing error event occurring at each particular site. Moreover we assume that (iii) if a site undergoes at least one mutation in the history of the sample, the site is segregating in the data; and (iv) any sequencing errors fall on each site with probability proportional to *P*. The validity of these assumptions is discussed below in Section S4.3.

The number of recurrent mutations in a sample with *m* variable sites can then be simulated using Algorithm 1. This is a ‘balls-into-bins’ type simulation, in which balls are placed one-by-one into *M* bins, each time selecting a bin at random with probability proportional to *P*, until *m* bins contain at least one ball; the output is the total number of balls thrown minus m. Executing Algorithm 1 multiple times and calculating a histogram of the results gives an approximation to the distribution of the number of recurrent mutations given the number *m* of observed segregating sites.

##### Algorithm 1: Simulating the number of recurrent mutations conditional on observing *m* variable sites

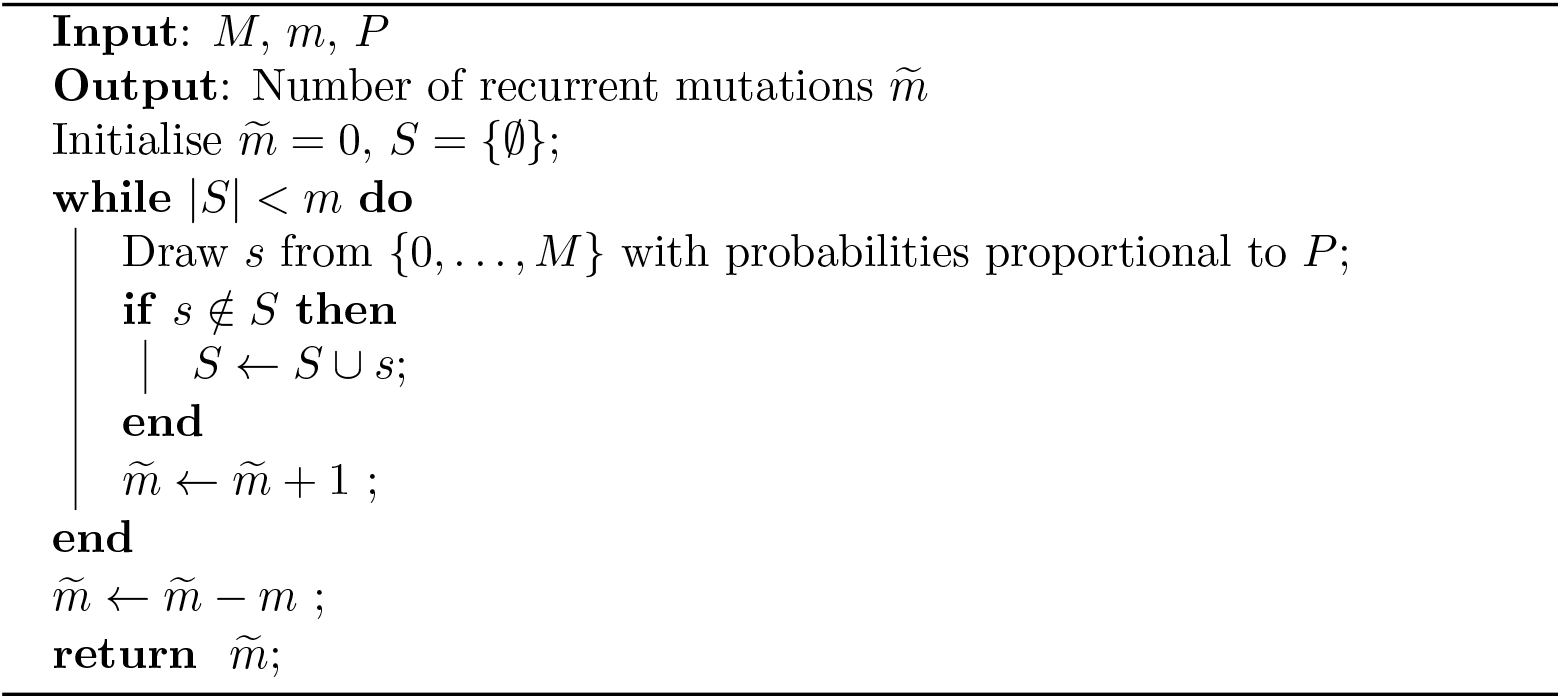

#### S4.2. Mutation rate heterogeneity along the genome

Parts of the genome with a relatively higher mutation rate are more likely to undergo recurrent mutation, so it is important to incorporate the effects of mutation rate heterogeneity. We use an empirical estimate of mutation density to approximate the variation in mutation rate along the genome.

##### S4.2.1. Data

All 17908 sequences in GISAID collected around the world between 1 and 3 February 2021 were downloaded, filtering for sequences labelled as complete (>29 000bp), high coverage, and excluding any with more than 5% ambiguous nucleotides. Alignment was performed as described in Section S1.1. SNP-sites (Page *et al*., 2016) was used to extract the positions of the 13 747 identified SNPs; a vector 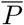 of length 29 903 was then formed, with a 1 entry at position *i* if there had been at least one mutation at position *i* of the genome, and 0 otherwise. If the mutation rate is constant along the genome, we would expect the 1’s to be spread uniformly throughout 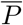; uneven clustering of the mutations gives an indication of mutation rate heterogeneity. We note that an alternative approach would be to fit a tree to the sequencing data (using maximum likelihood, for instance), count the minimum number of mutations required at each site of the genome, and use this to estimate *P*. However, this was found to result in very noisy estimates, and provide worse quantification of mutation rate heterogeneity (which we confirmed through simulation studies).

##### S4.2.2. Smoothing

The mutation density along the genome was then estimated nonparametrically from 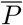 by smoothing using wavelet decomposition, as implemented in the R package `wavethresh` (Nason *et al*., 2010). This method was chosen as it does not require selecting a particular model, and it captures both fine-scale and broad variation in mutation density, allowing for the calculation of a smoothed estimate of 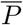 incorporating both local and large-scale rate heterogeneity.

Briefly, wavelet decomposition can be used to obtain an estimate of a signal from a set of discrete observations, by analysing variation in the data at increasingly coarser scales (Nason, 2008). Given *M* = 2^*n*^ observations of sites, corresponding to the entries of 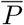 (padding the vector 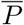 to the nearest power of 2 by reflecting the data at the endpoints), *n* iterations are performed, and at the *i*-th iteration (1) coefficients are computed using (non-overlapping) subsets of 2*^i^* neighbouring observations, and (2) these coefficients are used to refine a smoothed estimate of the data. The computation of coefficients and the smoothed approximations is governed by the choice of wavelet shape; we use Daubechies’ least-asymmetric wavelets (Daubechies, 1988) with six vanishing moments (other choices of wavelet basis produced similar results). Wavelet *shrinkage* can be used to obtain a smoothed estimate of the observations and remove noise: coefficient selection is performed by only keeping coefficients with values above a certain threshold and setting the others to zero. There are myriad ways of calculating such a threshold (Nason, 2008); we apply the empirical Bayes method of Johnstone & Silverman (2005b) implemented in the R package `EbayesThresh` (Johnstone & Silverman, 2005a).

As the mutation rate is dependent on the base type of the nucleotide undergoing mutation (Simmonds, 2020; Koyama *et al*., 2020), 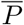 was split into four parts by the corresponding base type in the reference sequence, and the wavelet decomposition and thresholding performed separately for each part before joining them back together. The resulting smoothed estimate 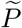 is shown in Figure 1. The total estimated mutation probability for each base type closely matches the actual proportion of mutations that fall on sites of each base type in the data, as desired. The smoothing method has clearly identified both localised and long-range variation in mutation density along the genome.

To check consistency of the results across time periods, data from September-November 2020 was also used to produce smoothed estimates of 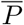 (consisting of 41376 sequences with 14 263 variable sites). The resulting estimate was found to agree closely to that obtained using the February data, so the latter was used in further analysis.

#### S4.3. Validity of assumptions

We now return to consider the validity of the assumptions stated in Section S4.1. Assumption (i) appears reasonable for the data at hand. Assumption (iii) can be violated if a mutation arising on a branch of the genealogy subsequently reverses through recurrent mutation: either on the same branch before it splits, or independently on every child branch subtending the mutation. We note that the probability of such events depends on the distribution of branch lengths in the genealogy; simulations using the standard coalescent model show that the probability of such events is small. Moreover, such events can never create incompatibilities in the data, so we can ignore their possibility for our purposes, as the solutions identified by KwARG will never include such recurrent mutation events.

Regarding assumption (ii), as the mutation rates depend on the base type, we cannot claim that all sequences have exactly the same probabilities *P* of mutating at each particular site, as this will depend on the nucleotides carried by the sequence. However, we estimate the effect of this violation to be negligible, given the relatively low overall rate of mutation for SARS-CoV-2.

To make our approximation even more conservative, we increase *m* by adding back the number of masked segregating sites (which are as stated in Sections S1.3 to S1.6), and further multiply the number of sites by a penalty factor of *F* = 1.1, which is justified in Section S4.3.1 below. Thus, we address assumption (iv) by noting that we have masked sites that are excessively prone to sequencing errors in the data, so correspondingly we decrease *M* by the number of masked sites and delete the corresponding entries from 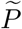. It is then reasonable to assume that sequencing errors occurring at the non-masked sites fall at each site with the same probabilities as mutations. The effects of this assumption being violated are explored further in Section S4.3.2.

##### S4.3.1. Choice of penalty factor

As noted above, the number of segregating sites in the sample is multiplied by a penalty factor *F* before performing the simulations. This results in a larger number of recurrent mutations being simulated, skewing the distribution to the right and thus ensuring that the *p*-values calculated from the simulated distribution are reasonably conservative. This is necessary because, as with any regression method that aims to (partially) de-noise the data, there is a risk that the fitted curve underestimates the true mutation rate heterogeneity, which would result in the expected number of recurrent mutations being underestimated, leading to false positives.

The choice of *F* = 1.1 was validated through simulation studies. First, we simulate a “true” mutation rate map *P*_true_ across 29,903 sites, as a realisation of an autoregressive process. Then, we simulate 20 000 mutations falling on the genome (allowing sites to mutate multiple times), and recreate the vector 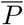 by marking which sites had (or had not) undergone at least one mutation. The method described in Section S4.2 is then applied to fit an estimated mutation density *P*_fit_. Finally, 10 000 simulations of Algorithm 1 are used to get an estimate of the null distribution: first, using *P*_true_ with *m* ∈ {100, 300, 500} sample segregating sites, then using *P*_sim_ with *m* · *F* sample segregating sites, for *F* ∈ {1.0, 1.1, 1.2, 1.3, 1.4, 1.5}.

This procedure was repeated 500 times for each combination of *m* and *F*. The results are presented in Figure S1. This demonstrates that without the penalty term, the fitted mutation density may indeed fail to capture all of the mutation rate heterogeneity that is present; for instance, when considering a sample with 300 segregating sites, in 46% of cases the 95th percentile of the simulated distribution will be lower than that of the true distribution. The results demonstrate that a value of *F* = 1.1 appears sufficient to negate this effect, without excessively increasing the false negative rate.

**Figure S1.**
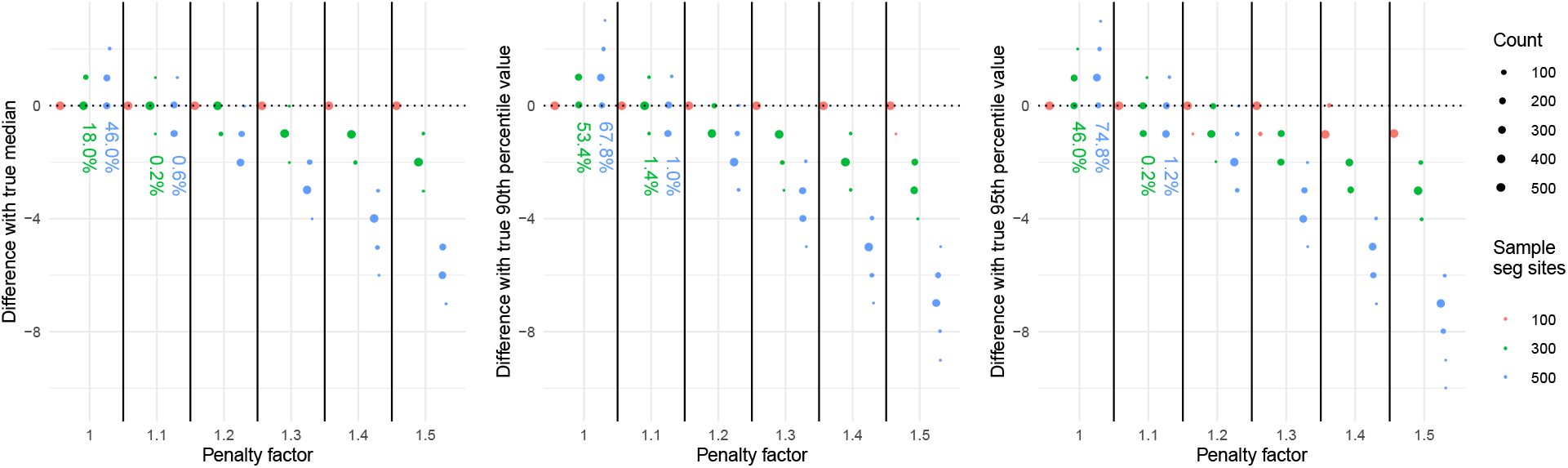
Comparison of simulated null distributions using *P*_true_ and *P*_fit_. Points show the difference between the true and simulated median (left panel), 90th percentile (middle), 95th percentile (right), with size proportional to the number of observations, split by penalty factor *F* (*x*-axis) and the number of sample segregating sites *m* (colours). Ideally, the points should be concentrated around 0; values above (below) 0 may result in false positives (false negatives) when using the estimated null distribution. Percentages show the proportion of cases lying above 0.

##### S4.3.2. Presence of highly homoplasic sites

Violations of assumption (iv) can occur if some (non-masked) sites along the genome are highly homoplasic, which can occur due to the effects of selection, or as an artifact of the sequencing process. Our method will not pick up the spikes in the corresponding positions of 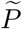, potentially introducing bias to the simulated null distribution that could lead to false positives.

We investigate the extent to which a violation of assumption (iv) affects the resulting inference through simulation studies. For each *i* ∈ {0,1, 5, 10, 20, 50, 100, 200}, *i* sites of the genome are chosen, and the corresponding probabilities in 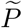 are multiplied by a factor *H* ∈ {2, 5, 10, 20, 50} to give the vectors 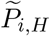. This recreates the effect of having i sites which are highly homoplasic (with the extent of this controlled by *H*); an example of 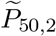 is shown in Figure S2.

For each combination of *i* and *H*, 200 datasets of 80 sequences were simulated using msprime (Kelleher *et al*., 2016), with parameters that appear reasonable for SARS-CoV-2:

**Figure S2.**
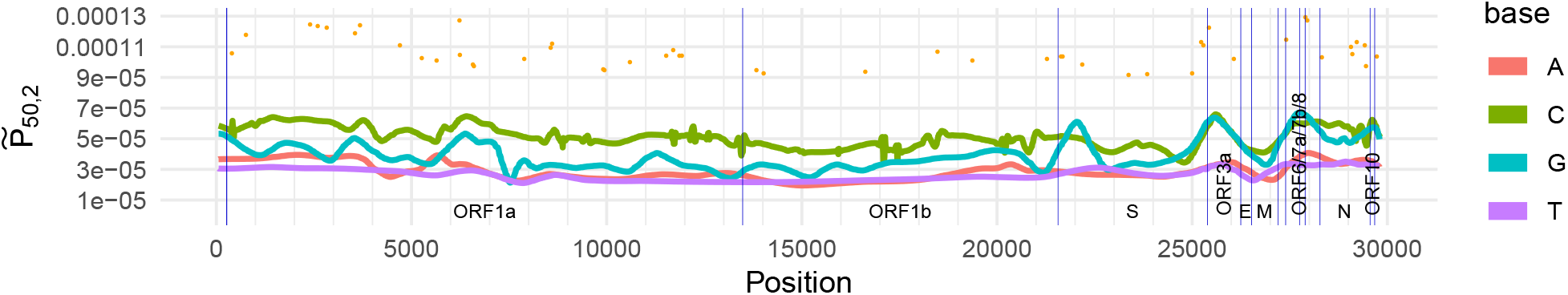
Mutation density estimate 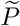 adjusted by selecting *i* = 50 sites and multiplying the corresponding entry of 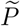 by *H* = 2 (resulting values shown in orange), to recreate the presence of 50 highly homoplasic sites.

- *N_e_* = 1 · 10^6^, exponential growth rate of 1.5 (no appropriate published estimates of these parameters could be identified, but this choice was found to give reasonable values of MRCA time and number of segregating sites for the simulated datasets);
- binary mutation model (finite sites);
- mutation rate per site per generation given by the entries of 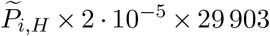. This was calculated based on:

- a mean mutation rate of 8 · 10^−4^ per site per year (as used by Nextstrain (Hadfield *et al*., 2018), accessed through nextstrain.org/ncov/global);
- a generation time of 7.5 days (Li *et al*., 2020);
- giving a mean mutation rate of 2 · 10^-5^ per site per generation.

For each dataset, KwARG was run 200 times (parameters: *T* = 30, *Q* = 100, (*C_SE_, C_RM_, C_R_, C_RR_*) ∈ {(1, 1.1, ∞, ∞), (0.01, 0.02, 1.00, 2.00)}) to calculate the minimal number of recurrent mutations needed to explain the dataset in the absence of recombination. A *p*-value was then calculated, using the null distribution simulated using the un-adjusted vector 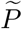 and 10 000 iterations of Algorithm 1, with *m* set to the number of segregating sites in the dataset multiplied by the penalty factor *F* = 1.1.

The proportion of times the null hypothesis is (incorrectly) rejected, with *p* < 0.05, is shown in the left panel of Figure 2.

##### S4.3.3. Detection rate vs recombination rate

We investigate how the proportion of cases in which the null hypothesis is rejected (with *p* < 0.05) varies with recombination rate. For several values of the recombination rate 1 · 10^−7^ ≤ *ρ* ≤ 1 · 10^−5^ (per site per generation), 200 datasets were simulated using msprime with the parameters given in Section S4.3.2 (using the un-adjusted vector 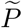), and the same method used to calculate a *p*-value for each dataset. The results are presented in the right panel of Figure 2.

### S5. Null distribution for MERS-CoV

The same methodology as described in Section S4 was used to simulate the null distribution for MERS-CoV.

**Figure S3.**
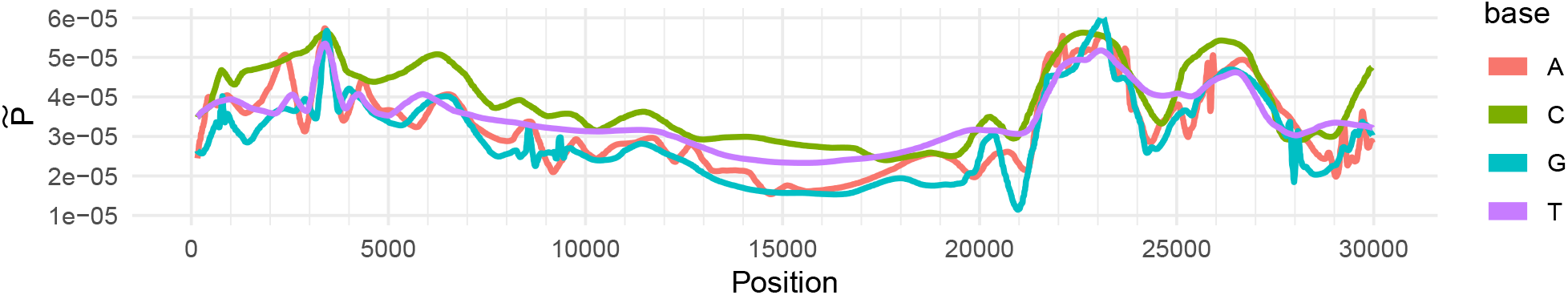
Estimate 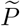 of the probability of a mutation falling on each site of the MERS-CoV genome.

Sequences were downloaded from the NCBI Virus database, filtering for those of length at least 20 000 bp, from human and camel hosts, across all time periods. Alignment to the reference sequence was performed as described in Section S2. The alignment comprised 700 sequences with 14 238 variable sites. The vector 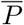 was constructed, and wavelet decomposition was used to fit the estimate 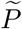 in the same manner as described in Section S4.2; the result is shown in Figure S3.

With the resulting estimate 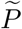 and the parameters given in Section S6, 1000 000 iterations of Algorithm 1 were used to simulate the null distribution. The resulting probabilities and *p*-values are shown in the third and fourth columns of Table 1.

### S6. Null distribution simulation parameters

|  | SA<br>(November) | SA<br>(February) | England<br>(November) | England<br>(January) | MERS-CoV |
| --- | --- | --- | --- | --- | --- |
| No. of segregating sites | 206 | 150 | 363 | 276 | 197 |
| Plus masked sites | 29 | 18 | 10 | 35 | 2 |
| Times penalty factor $F$ | 1.1 | 1.1 | 1.1 | 1.1 | 1.1 |
| $m$ | 259 | 185 | 410 | 342 | 219 |
| Length of genome | 29 903 | 29 903 | 29 903 | 29 903 | 30 119 |
| Less number of masked sites | 1126 | 1126 | 477 | 660 | 300 |
| $M$ (= length of $\tilde{P}$) | 28 777 | 28 777 | 29 426 | 29 243 | 29 819 |

---

**Table S1.** GISAID accession numbers, collection dates, and references of sequences in the South Africa (November) sample.

| Other lineages |  |  | Lineage B.1.351 |  |  |
| --- | --- | --- | --- | --- | --- |
| Accession | Date | Ref | Accession | Date | Ref |
| EPI_ISL_660225 | 02/11/2020 | SAO1 | EPI_ISL_736958 | 20/11/2020 | SAN1 |
| EPI_ISL_660257 | 18/11/2020 | SAO2 | EPI_ISL_696481 | 19/11/2020 | SAN2 |
| EPI_ISL_736993 | 25/11/2020 | SAO3 | EPI_ISL_660637 | 03/11/2020 | SAN3 |
| EPI_ISL_660643 | 01/11/2020 | SAO4 | EPI_ISL_678632 | 11/11/2020 | SAN4 |
| EPI_ISL_660229 | 16/11/2020 | SAO5 | EPI_ISL_736932 | 25/11/2020 | SAN5 |
| EPI_ISL_736985 | 25/11/2020 | SAO6 | EPI_ISL_678641 | 12/11/2020 | SAN6 |
| EPI_ISL_736926 | 26/11/2020 | SAO7 | EPI_ISL_700422 | 04/11/2020 | SAN7 |
| EPI_ISL_696462 | 19/11/2020 | SAO8 | EPI_ISL_696503 | 25/11/2020 | SAN8 |
| EPI_ISL_660655 | 03/11/2020 | SAO9 | EPI_ISL_700470 | 12/11/2020 | SAN9 |
| EPI_ISL_660625 | 05/11/2020 | SAO10 | EPI_ISL_736983 | 24/11/2020 | SAN10 |
| EPI_ISL_660231 | 16/11/2020 | SAO11 | EPI_ISL_736936 | 19/11/2020 | SAN11 |
| EPI_ISL_678608 | 15/11/2020 | SAO12 | EPI_ISL_700487 | 06/11/2020 | SAN12 |
| EPI_ISL_660163 | 05/11/2020 | SAO13 | EPI_ISL_736935 | 26/11/2020 | SAN13 |
| EPI_ISL_660232 | 17/11/2020 | SAO14 | EPI_ISL_700443 | 13/11/2020 | SAN14 |
| EPI_ISL_700488 | 05/11/2020 | SAO15 | EPI_ISL_736939 | 24/11/2020 | SAN15 |
| EPI_ISL_660652 | 01/11/2020 | SAO16 | EPI_ISL_700554 | 02/11/2020 | SAN16 |
| EPI_ISL_660622 | 07/11/2020 | SAO17 | EPI_ISL_696505 | 25/11/2020 | SAN17 |
| EPI_ISL_660651 | 02/11/2020 | SAO18 | EPI_ISL_696518 | 24/11/2020 | SAN18 |
| EPI_ISL_678612 | 15/11/2020 | SAO19 | EPI_ISL_700589 | 12/11/2020 | SAN19 |
| EPI_ISL_696509 | 24/11/2020 | SAO20 | EPI_ISL_736959 | 20/11/2020 | SAN20 |
| EPI_ISL_678595 | 18/11/2020 | SAO21 | EPI_ISL_696453 | 20/11/2020 | SAN21 |
| EPI_ISL_660222 | 09/11/2020 | SAO22 | EPI_ISL_696521 | 24/11/2020 | SAN22 |
| EPI_ISL_696468 | 18/11/2020 | SAO23 | EPI_ISL_736964 | 19/11/2020 | SAN23 |
| EPI_ISL_660230 | 16/11/2020 | SAO24 | EPI_ISL_736928 | 24/11/2020 | SAN24 |
| EPI_ISL_660626 | 07/11/2020 | SAO25 | EPI_ISL_678629 | 13/11/2020 | SAN25 |

**Table S2.** GISAID accession numbers, collection dates, and references of sequences in the South Africa (February) sample.

| Accession | Date | Accession | Date |
| --- | --- | --- | --- |
| EPI_ISL_1048548 | 01/02/2021 | EPI_ISL_1371925 | 15/02/2021 |
| EPI_ISL_1048553 | 02/02/2021 | EPI_ISL_1371926 | 09/02/2021 |
| EPI_ISL_1048554 | 02/02/2021 | EPI_ISL_1371927 | 19/02/2021 |
| EPI_ISL_1048555 | 02/02/2021 | EPI_ISL_1371928 | 21/02/2021 |
| EPI_ISL_1048562 | 01/02/2021 | EPI_ISL_1371929 | 20/02/2021 |
| EPI_ISL_1366778 | 02/02/2021 | EPI_ISL_1371930 | 09/02/2021 |
| EPI_ISL_1366779 | 02/02/2021 | EPI_ISL_1371931 | 21/02/2021 |
| EPI_ISL_1366781 | 01/02/2021 | EPI_ISL_1371932 | 17/02/2021 |
| EPI_ISL_1366782 | 18/02/2021 | EPI_ISL_1371933 | 23/02/2021 |
| EPI_ISL_1366783 | 25/02/2021 | EPI_ISL_1371995 | 05/02/2021 |
| EPI_ISL_1366793 | 04/02/2021 | EPI_ISL_1371996 | 15/02/2021 |
| EPI_ISL_1366840 | 05/02/2021 | EPI_ISL_1371999 | 09/02/2021 |
| EPI_ISL_1366864 | 05/02/2021 | EPI_ISL_1372000 | 07/02/2021 |
| EPI_ISL_1366869 | 05/02/2021 | EPI_ISL_1372001 | 08/02/2021 |
| EPI_ISL_1366877 | 05/02/2021 | EPI_ISL_1372002 | 09/02/2021 |
| EPI_ISL_1366887 | 06/02/2021 | EPI_ISL_1372003 | 24/02/2021 |
| EPI_ISL_1366888 | 05/02/2021 | EPI_ISL_1372004 | 18/02/2021 |
| EPI_ISL_1371923 | 23/02/2021 | EPI_ISL_1372005 | 17/02/2021 |
| EPI_ISL_1371924 | 21/02/2021 | EPI_ISL_1372006 | 17/02/2021 |

**Table S3.** GISAID accession numbers, collection dates, and references of sequences in the England (November) sample.

| Other lineages |  |  | Lineage B.1.1.7 |  |  |
| --- | --- | --- | --- | --- | --- |
| Accession | Date | Ref | Accession | Date | Ref |
| EPI_ISL_662468 | 12/11/2020 | EO1 | EPI_ISL_708881 | 30/11/2020 | EN1 |
| EPI_ISL_664402 | 06/11/2020 | EO2 | EPI_ISL_705071 | 22/11/2020 | EN2 |
| EPI_ISL_702752 | 19/11/2020 | EO3 | EPI_ISL_657548 | 09/11/2020 | EN3 |
| EPI_ISL_650455 | 13/11/2020 | EO4 | EPI_ISL_702338 | 27/11/2020 | EN4 |
| EPI_ISL_667977 | 14/11/2020 | EO5 | EPI_ISL_656730 | 08/11/2020 | EN5 |
| EPI_ISL_642566 | 02/11/2020 | EO6 | EPI_ISL_709730 | 26/11/2020 | EN6 |
| EPI_ISL_661404 | 11/11/2020 | EO7 | EPI_ISL_702093 | 28/11/2020 | EN7 |
| EPI_ISL_679726 | 01/11/2020 | EO8 | EPI_ISL_675080 | 15/11/2020 | EN8 |
| EPI_ISL_654967 | 10/11/2020 | EO9 | EPI_ISL_673518 | 15/11/2020 | EN9 |
| EPI_ISL_659205 | 05/11/2020 | EO10 | EPI_ISL_704716 | 30/11/2020 | EN10 |
| EPI_ISL_659013 | 01/11/2020 | EO11 | EPI_ISL_676036 | 13/11/2020 | EN11 |
| EPI_ISL_662253 | 11/11/2020 | EO12 | EPI_ISL_704695 | 02/11/2020 | EN12 |
| EPI_ISL_660027 | 04/11/2020 | EO13 | EPI_ISL_704619 | 21/11/2020 | EN13 |
| EPI_ISL_646293 | 04/11/2020 | EO14 | EPI_ISL_658341 | 08/11/2020 | EN14 |
| EPI_ISL_664758 | 12/11/2020 | EO15 | EPI_ISL_661750 | 14/11/2020 | EN15 |
| EPI_ISL_659140 | 05/11/2020 | EO16 | EPI_ISL_665414 | 02/11/2020 | EN16 |
| EPI_ISL_661929 | 14/11/2020 | EO17 | EPI_ISL_703736 | 26/11/2020 | EN17 |
| EPI_ISL_641906 | 03/11/2020 | EO18 | EPI_ISL_658292 | 08/11/2020 | EN18 |
| EPI_ISL_661483 | 11/11/2020 | EO19 | EPI_ISL_709568 | 26/11/2020 | EN19 |
| EPI_ISL_656165 | 06/11/2020 | EO20 | EPI_ISL_704601 | 22/11/2020 | EN20 |
| EPI_ISL_658415 | 08/11/2020 | EO21 | EPI_ISL_656409 | 08/11/2020 | EN21 |
| EPI_ISL_655916 | 08/11/2020 | EO22 | EPI_ISL_668252 | 12/11/2020 | EN22 |
| EPI_ISL_637180 | 02/11/2020 | EO23 | EPI_ISL_661854 | 12/11/2020 | EN23 |
| EPI_ISL_673482 | 15/11/2020 | EO24 | EPI_ISL_703229 | 19/11/2020 | EN24 |
| EPI_ISL_703087 | 19/11/2020 | EO25 | EPI_ISL_657799 | 08/11/2020 | EN25 |
| EPI_ISL_675115 | 13/11/2020 | EO26 | EPI_ISL_708945 | 30/11/2020 | EN26 |
| EPI_ISL_664943 | 04/11/2020 | EO27 | EPI_ISL_679428 | 22/11/2020 | EN27 |
| EPI_ISL_706068 | 02/11/2020 | EO28 | EPI_ISL_676194 | 13/11/2020 | EN28 |
| EPI_ISL_657282 | 08/11/2020 | EO29 | EPI_ISL_683471 | 24/11/2020 | EN29 |
| EPI_ISL_679916 | 06/11/2020 | EO30 | EPI_ISL_676012 | 13/11/2020 | EN30 |
| EPI_ISL_673815 | 15/11/2020 | EO31 | EPI_ISL_705063 | 22/11/2020 | EN31 |
| EPI_ISL_678719 | 16/11/2020 | EO32 | EPI_ISL_659491 | 05/11/2020 | EN32 |
| EPI_ISL_705061 | 19/11/2020 | EO33 | EPI_ISL_668018 | 12/11/2020 | EN33 |
| EPI_ISL_646457 | 03/11/2020 | EO34 | EPI_ISL_702918 | 19/11/2020 | EN34 |
| EPI_ISL_656970 | 08/11/2020 | EO35 | EPI_ISL_657622 | 08/11/2020 | EN35 |
| EPI_ISL_647347 | 01/11/2020 | EO36 | EPI_ISL_704698 | 01/11/2020 | EN36 |
| EPI_ISL_650406 | 08/11/2020 | EO37 | EPI_ISL_679302 | 21/11/2020 | EN37 |
| EPI_ISL_661700 | 13/11/2020 | EO38 | EPI_ISL_704606 | 22/11/2020 | EN38 |
| EPI_ISL_658474 | 08/11/2020 | EO39 | EPI_ISL_703148 | 19/11/2020 | EN39 |
| EPI_ISL_700654 | 09/11/2020 | EO40 | EPI_ISL_645527 | 05/11/2020 | EN40 |

**Table S4.**
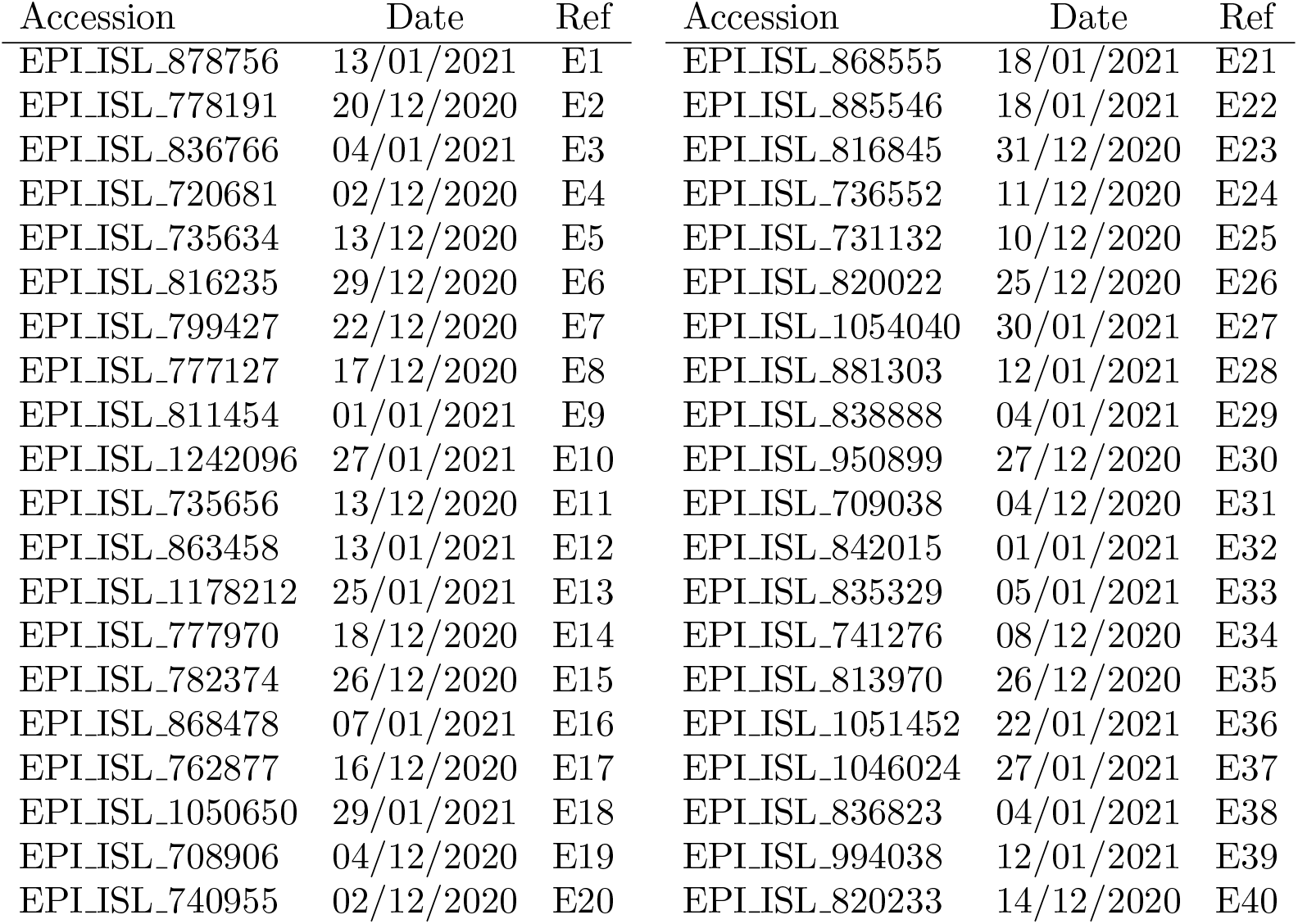
GISAID accession numbers, collection dates, and references of sequences in the England (January) sample.

**Table S5.** NCBI Virus database accession numbers, collection dates, and references of sequences in the MERS-CoV sample.

| Accession | Submitters | Date | Ref |
| --- | --- | --- | --- |
| KY688118.1 | Paden, C. R., et al. | 07/02/2015 | M1 |
| KT806044.1 | Lu, X., et al. | 09/02/2015 | M2 |
| KT806045.1 | Lu, X., et al. | 22/02/2015 | M3 |
| KT806047.1 | Lu, X., et al. | 27/03/2015 | M4 |
| KT806048.1 | Lu, X., et al. | 07/02/2015 | M5 |
| KT806049.1 | Lu, X., et al. | 15/02/2015 | M6 |
| KT806051.1 | Lu, X., et al. | 05/02/2015 | M7 |
| KT806052.1 | Lu, X., et al. | 02/02/2015 | M8 |
| KT806053.1 | Lu, X., et al. | 02/02/2015 | M9 |
| KT806054.1 | Lu, X., et al. | 13/02/2015 | M10 |
| KT806055.1 | Lu, X., et al. | 10/02/2015 | M11 |
| KT026453.1 | Park, W. B., et al. | 10/02/2015 | M12 |
| KT026454.1 | Park, W. B., et al. | 01/03/2015 | M13 |
| KT026455.1 | Park, W. B., et al. | 10/02/2015 | M14 |
| KT026456.1 | Park, W. B., et al. | 01/03/2015 | M15 |
| KR011263.1 | Lu, X., et al. | 21/01/2015 | M16 |
| KR011264.1 | Lu, X., et al. | 21/01/2015 | M17 |
| KR011265.1 | Lu, X., et al. | 26/01/2015 | M18 |
| KR011266.1 | Lu, X., et al. | 06/01/2015 | M19 |

**Figure S4.**
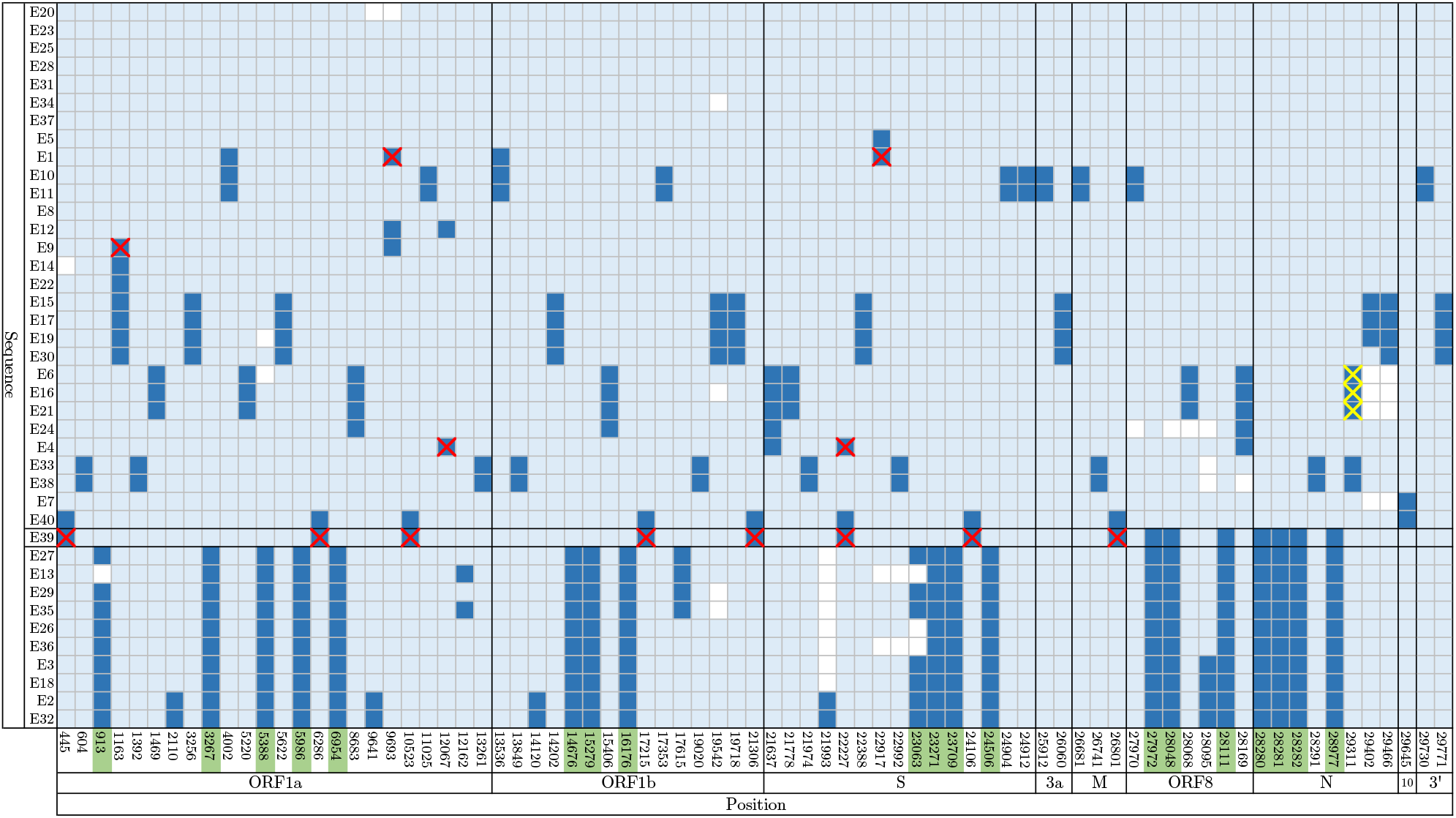
Summary of the England (January) dataset. Rows correspond to sequences, labelled on the left. Columns correspond to positions along the genome; uninformative sites (with all 0’s or 1’s) and those with singleton mutations (with exactly one 1) are not shown. Light blue: ancestral state, dark blue: mutated state, white: missing data. Red crosses highlight sites of recurrent mutations identified by KwARG located on the terminal branches of the ARG (affecting only one sequence). Yellow crosses highlight recurrent mutations on internal branches (hence affecting multiple sequences). Sites bearing the characteristic mutations of lineage B.1.1.7 (Rambaut *et al*., 2020) are highlighted in green.

**Figure S5.**
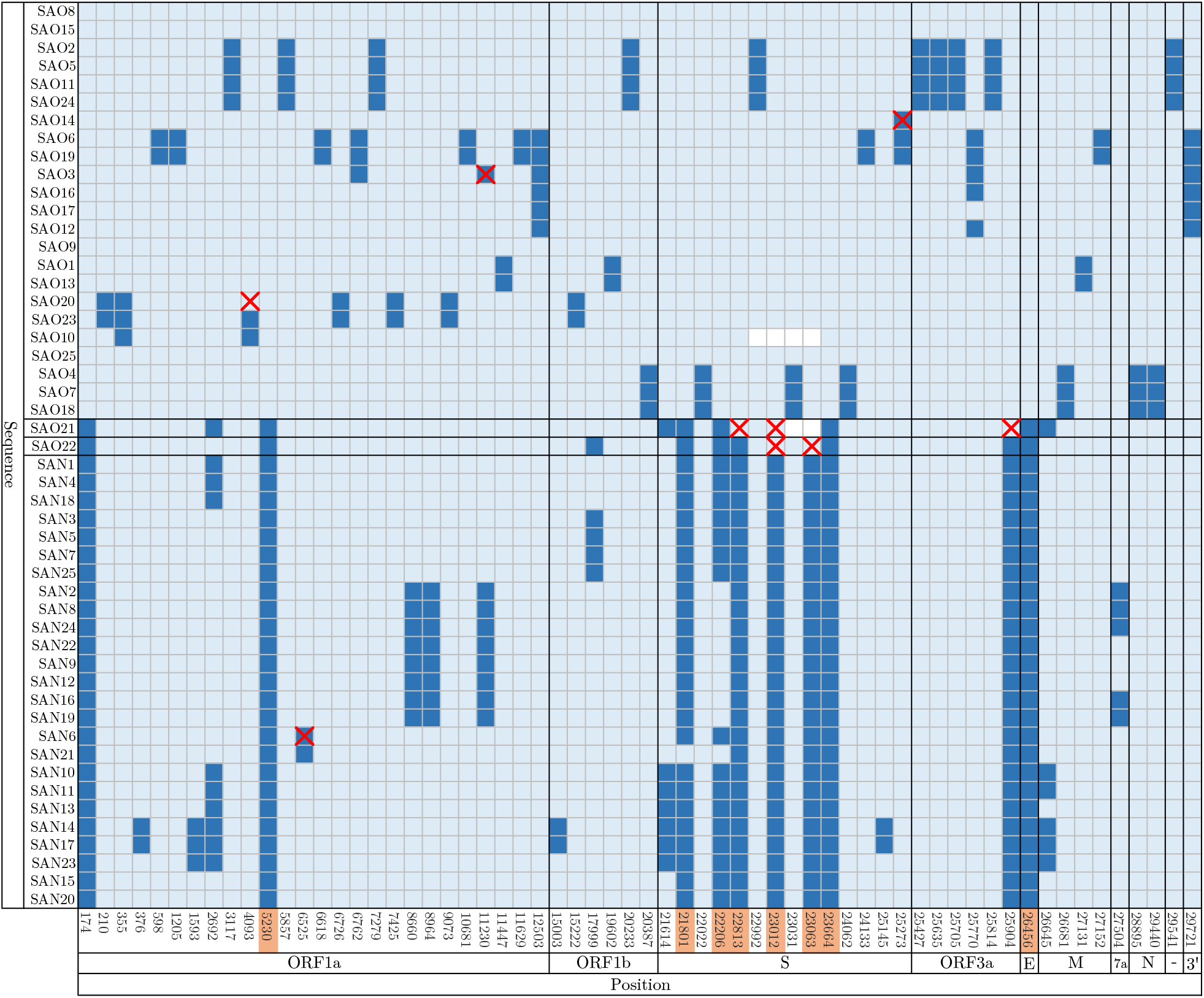
Summary of the South Africa (November) dataset. Rows correspond to sequences, labelled on the left. Columns correspond to positions along the genome; uninformative sites (with all 0’s or 1’s) and those with singleton mutations (with exactly one 1) are not shown. Light blue: ancestral state, dark blue: mutated state, white: missing data. Red crosses highlight sites of recurrent mutations identified by KwARG. Sites bearing the characteristic (non-synonymous) mutations of lineage B.1.351 (Tegally *et al*., 2020) are highlighted in orange.

**Figure S6.**
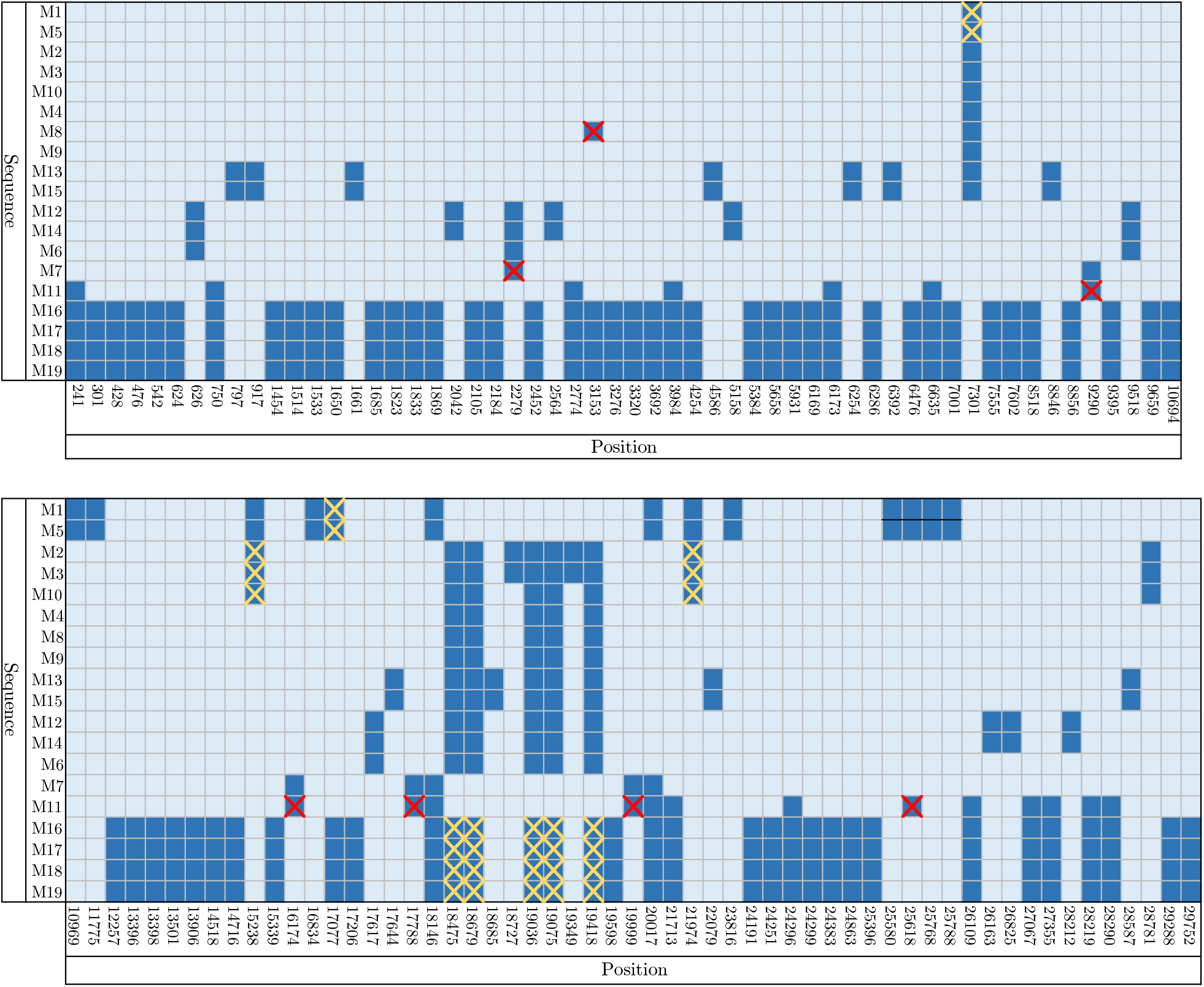
Summary of the MERS dataset. Rows correspond to sequences, labelled on the left. Columns correspond to positions along the genome; uninformative sites (with all 0’s or 1’s) and those with singleton mutations (with exactly one 0 or 1) are not shown. Light blue and dark blue denote differing allele types. Red crosses highlight sites of recurrent mutations identified by KwARG located on the terminal branches of the ARG (affecting only one sequence). Yellow crosses highlight recurrent mutations on internal branches (hence affecting multiple sequences).

**Figure S7.**
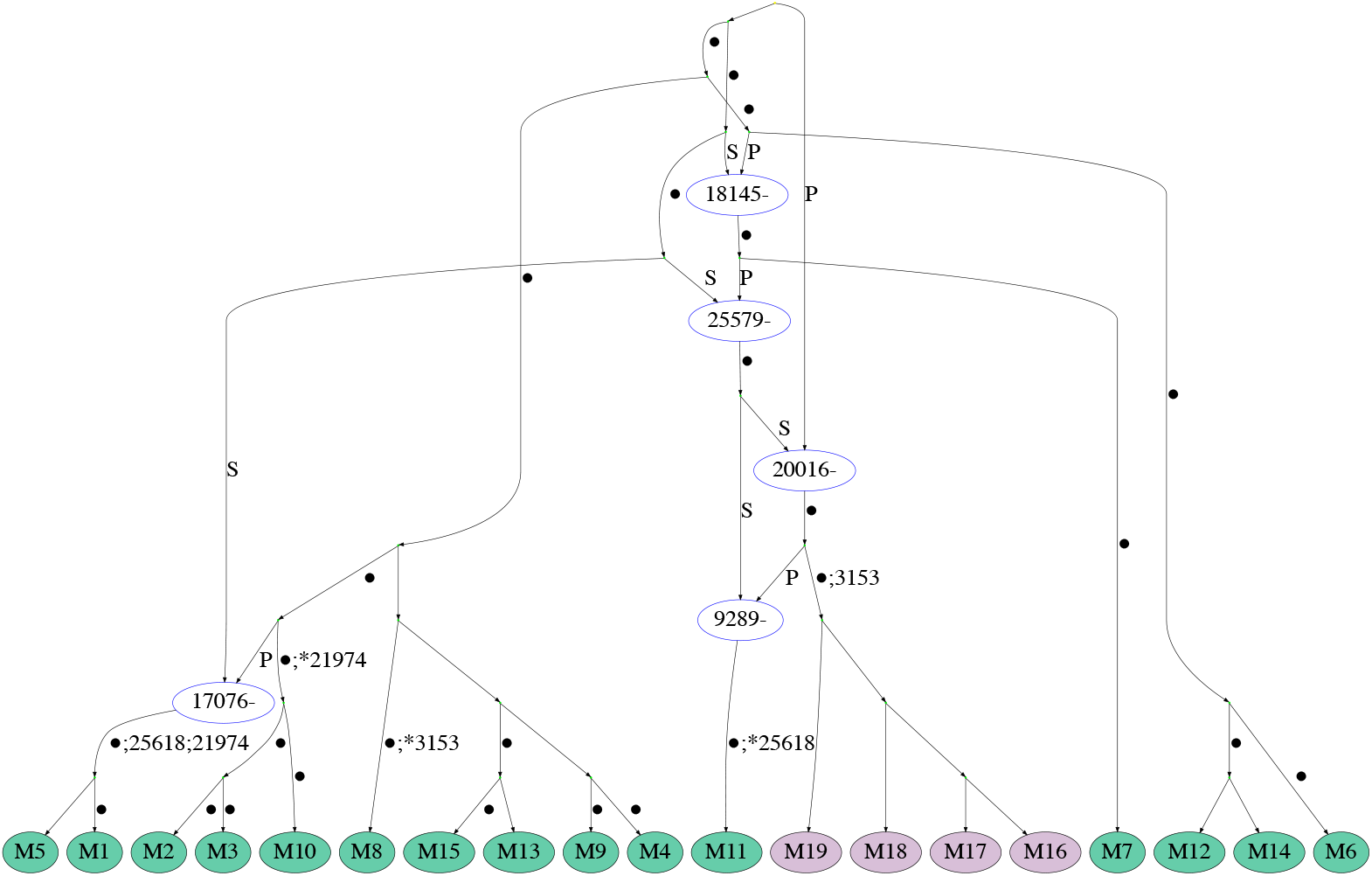
Example of an ARG for the MERS-CoV dataset. Recombination nodes are shown in blue, labelled with the recombination breakpoint, with the offspring sequence inheriting part of the genome to the left (right) of the breakpoint from the parent labelled “P” (“S”). Recurrent mutations are prefixed with an asterisk. Edges are labelled by positions of mutations (some mutated sites are not explicitly labelled and are denoted by a dot instead).

## References

1. De Maio, N. et al. Issues with SARS-CoV-2 sequencing data https://virological.org/t/issues-with-sars-cov-2-sequencing-data/473. 2020.

2. Dudas, G. & Rambaut, A. MERS-CoV recombination: implications about the reservoir and potential for adaptation. Virus Evolution 2(2016).

3. Elbe, S. & Buckland-Merrett, G. Data, disease and diplomacy: GISAID’s innovative contribution to global health. Global Challenges 1, 33–46 (2017).

4. Gribble, J. et al. The coronavirus proofreading exoribonuclease mediates extensive viral recombination. PLOS Pathogens 17, e1009226 (2021).

5. Griffiths, R. C. & Marjoram, P. *An ancestral recombination graph* in Progress in population genetics and human evolution (Springer, 1997), 257–270.

6. Hatcher, E. L. et al. Virus Variation Resource–improved response to emergent viral outbreaks. Nucleic Acids Research 45, D482–D490 (2017).

7. Hein, J., Schierup, M. & Wiuf, C. Gene genealogies, variation and evolution: A primer in coalescent theory (Oxford University Press, USA, 2004).

8. Hudson, R. R. & Kaplan, N. L. Statistical properties of the number of recombination events in the history of a sample of DNA sequences. Genetics 111, 147–164 (1985).

9. Ignatieva, A., Lyngsø, R. B., Jenkins, P. A. & Hein, J. KwARG: Parsimonious reconstruction of ancestral recombination graphs with recurrent mutation. Bioinformatics, in press (2021).

10. Jackson, B. et al. Recombinant SARS-CoV-2 genomes involving lineage B.1.1.7 in the UK https://virological.org/t/recombinant-sars-cov-2-genomes-involving-lineage-b-1-1-7-in-the-uk/658. 2021.

11. Koyama, T., Platt, D. & Parida, L. Variant analysis of SARS-CoV-2 genomes. Bulletin of the World Health Organization 98, 495 (2020).

12. Maynard Smith, J & Smith, N. H. Detecting recombination from gene trees. Molecular Biology and Evolution 15, 590–599 (1998).

13. McVean, G., Awadalla, P. & Fearnhead, P. A coalescent-based method for detecting and estimating recombination from gene sequences. Genetics 160, 1231–1241 (2002).

14. Nie, Q. et al. Phylogenetic and phylodynamic analyses of SARS-CoV-2. Virus Research 287, 198098 (2020).

15. Posada, D. & Crandall, K. A. Evaluation of methods for detecting recombination from DNA sequences: Computer simulations. PNAS 98, 13757–13762 (2001).

16. Posada, D. & Crandall, K. A. The effect of recombination on the accuracy of phylogeny estimation. Journal of Molecular Evolution 54, 396–402 (2002).

17. Rambaut, A et al. Preliminary genomic characterisation of an emergent SARS-CoV-2 lineage in the UK defined by a novel set of spike mutations https://virological.org/t/preliminary-genomic-characterisation-of-an-emergent-sars-cov-2-lineage-in-the-uk-defined-by-a-novel-set-of-spike-mutations/563. 2020.

18. Richard, D., Owen, C. J., van Dorp, L. & Balloux, F. No detectable signal for ongoing genetic recombination in SARS-CoV-2. bioRxiv. doi:10.1101/2020.12.15.422866 (2020).

19. Sabir, J. S. et al. Co-circulation of three camel coronavirus species and recombination of MERS-CoVs in Saudi Arabia. Science 351, 81–84 (2016).

20. Samoilov, A. et al. Change of dominant strain during dual SARS-CoV-2 infection. medRxiv.doi:10.1101/2020.11.29.20238402 (2020).

21. Schierup, M. H. & Hein, J. Consequences of recombination on traditional phylogenetic analysis. Genetics 156, 879–891 (2000).

22. Simmonds, P. Rampant C→U hypermutation in the genomes of SARS-CoV-2 and other coronaviruses: Causes and consequences for their short- and long-term evolutionary trajectories. mSphere 5(2020).

23. Simon-Loriere, E. & Holmes, E. C. Why do RNA viruses recombine? Nature Reviews Microbiology 9, 617–626 (2011).

24. Su, S. et al. Epidemiology, genetic recombination, and pathogenesis of coronaviruses. Trends in Microbiology 24, 490–502 (2016).

25. Tang, X. et al. On the origin and continuing evolution of SARS-CoV-2. National Science Review 7, 1012–1023 (2020).

26. Tegally, H. et al. Emergence and rapid spread of a new severe acute respiratory syndrome-related coronavirus 2 (SARS-CoV-2) lineage with multiple spike mutations in South Africa. medRxiv.doi:10.1101/2020.12.21.20248640 (2020).

27. van Dorp, L. et al. Emergence of genomic diversity and recurrent mutations in SARS-CoV-2. Infection, Genetics and Evolution 83, 104351 (2020).

28. van Dorp, L. et al. No evidence for increased transmissibility from recurrent mutations in SARS-CoV-2. Nature Communications 11 (2020).

29. VanInsberghe, D., Neish, A., Lowen, A. C. & Koelle, K. Identification of SARS-CoV-2 recombinant genomes. bioRxiv. doi: 10.1101/2020.08.05.238386 (2020).

30. Varabyou, A., Pockrandt, C., Salzberg, S. L. & Pertea, M. Rapid detection of inter-clade recombination in SARS-CoV-2 with Bolotie. bioRxiv. doi:10.1101/2020.09.21.300913 (2020).

31. Wang, H., Kosakovsky Pond, S. L., Nekrutenko, A. & Nielsen, R. Testing recombination in the pandemic SARS-CoV-2 strains https://virological.org/t/testing-recombination-in-the-pandemic-sars-cov-2-strains/492. 2020.

32. Wu, F. et al. A new coronavirus associated with human respiratory disease in China. Nature 579, 265–269 (2020).

33. Yi, H. 2019 novel coronavirus is undergoing active recombination. Clinical Infectious Diseases 71, 884–887 (2020).

34. Zhang, Z., Shen, L. & Gu, X. Evolutionary dynamics of MERS-CoV: potential recombination, positive selection and transmission. Scientific Reports 6, 1–10 (2016).

## References

1. Daubechies, I. Orthonormal bases of compactly supported wavelets. Communications on Pure and Applied Mathematics 41, 909–996 (1988).

2. De Maio, N. et al. Issues with SARS-CoV-2 sequencing data https://virological.org/t/issues-with-sars-cov-2-sequencing-data/473. 2020.

4. Hadfield, J. et al. Nextstrain: Real-time tracking of pathogen evolution. Bioinformatics 34, 4121–4123 (2018).

5. Hatcher, E. L. et al. Virus Variation Resource-improved response to emergent viral outbreaks. Nucleic Acids Research 45, D482–D490 (2017).

6. Jackson, B. et al. Recombinant SARS-CoV-2 genomes involving lineage B.1.1.7 in the UK https://virological.org/t/recombinant-sars-cov-2-genomes-involving-lineage-b-1-1-7-in-the-uk/658. 2021.

7. Johnstone, I. M. & Silverman, B. W. EbayesThresh: R and S-Plus programs for empirical Bayes thresholding. Journal of Statistical Software 12, 1–38 (2005).

8. Johnstone, I. M. & Silverman, B. W. Empirical Bayes selection of wavelet thresholds. Annals of Statistics, 1700–1752 (2005).

9. Katoh, K. & Standley, D. M. MAFFT multiple sequence alignment software version 7: Improvements in performance and usability. Molecular Biology and Evolution 30, 772–780 (2013).

10. Kelleher, J., Etheridge, A. M. & McVean, G. Efficient coalescent simulation and genealogical analysis for large sample sizes. PLoS Computational Biology 12, e1004842 (2016).

12. Li, Q. et al. Early transmission dynamics in Wuhan, China, of novel coronavirus-infected pneumonia. New England Journal of Medicine (2020).

13. Nason, G. Wavelet methods in statistics with R (Springer Science & Business Media, 2008).

14. Nason, G. et al. Wavethresh: Wavelets statistics and transforms, v.4.6.8 https://CRAN.R-project.org/package=wavethresh. 2010.

15. Page, A. J. et al. SNP-sites: Rapid efficient extraction of SNPs from multi-FASTA alignments. Microbial Genomics 2 (2016).

16. Shen, W., Le, S., Li, Y. & Hu, F. SeqKit: A cross-platform and ultrafast toolkit for FASTA/Q file manipulation. PLOS ONE 11, e0163962 (2016).

17. Simmonds, P. Rampant C→U hypermutation in the genomes of SARS-CoV-2 and other coronaviruses: Causes and consequences for their short- and long-term evolutionary trajectories. mSphere 5(2020).

18. van Dorp, L. et al. Emergence of genomic diversity and recurrent mutations in SARS-CoV-2. Infection, Genetics and Evolution 83, 104351 (2020).

19. Wu, F. et al. A new coronavirus associated with human respiratory disease in China. Nature 579, 265–269 (2020).

